# Comprehensive classification of HCN1 variants linked to neurodevelopmental disorders with and without epilepsy

**DOI:** 10.64898/2026.03.18.712601

**Authors:** Roberta Castelli, Carla Marini, Alessandro Porro, Anna Castellini, Greta Fontana, Andrea Saponaro, Gianpiero Cavalleri, Susanna Rizzi, Carlo Fusco, Amitav Parida, Richard Caswell, Charlotte Sherlaw, Dario Pruna, Chris Reid, Lauren E. Bleakley, Katherine B. Howell, Ingrid Sheffer, Vishnu Anad Cuddapah, Shimriet Zeidler, Elena Pavlidis, Deb Pal, Krzysztof Szczałuba, Ghayda Mirzaa, Nathalie Couque, Yline Capri, Laurence Faivre, Frederic Tran Mau-Them, Fabio Sirchia, Christian M. Korff, Dario DiFrancesco, Gerhard Thiel, Christel Depienne, Bina Santoro, Anna Moroni

**Author notes:** Correspondence to: Anna Moroni, Department of Biosciences, University of Milan, via Celoria 26, 20133, Milan, Italy.

## Abstract

Hyperpolarization-activated cyclic nucleotide-gated 1 channels (HCN1) mediate the I*_h_* cationic current and play a central role in regulating neuronal excitability and synaptic integration. HCN1 is predominantly expressed in the neocortex and hippocampus. Pathogenic variants in *HCN1* have been increasingly identified in individuals presenting with a broad spectrum of epileptic disorders, ranging from severe developmental and epileptic encephalopathy (DEE) to milder epilepsies. Here, we used patch-clamp electrophysiology in combination with confocal imaging in HEK293 cells to functionally characterize 43 *HCN1* variants found in patients presenting with neurodevelopmental disorders, with or without epilepsy. Based on their biophysical properties, we defined four functional classes: (I) low or no current, (II) hyperpolarizing (i.e. left) shift in voltage dependence, (III) depolarizing (i.e. right) shift in voltage dependence, and (IV) generation of an instantaneous current. Integration of this functional classification with detailed clinical data from a cohort of 49 patients revealed a striking genotype-phenotype correlation. Loss-of-function variants were strongly enriched among individuals without epilepsy or with milder generalized phenotypes, whereas gain-of-function and mixed variants were predominantly associated with epilepsy, including all cases of DEE. Notably, non-epileptic cases clustered within a subgroup of loss-of-function variants affecting the selectivity filter. We further show that allosteric modulators, including the peptides NB6 and TRIP8b_nano_ and the small molecule J&J12e, normalize the functional properties of mutant HCN1 channels in three classes. These findings establish a clinically relevant framework for interpreting *HCN1* gain- and loss-of-function variants suggesting that the direction of channel dysfunction is a major determinant of epilepsy risk and severity.

## Introduction

Hyperpolarization-activated cyclic nucleotide-gated (HCN) channels are a family of four membrane proteins, encoded by the *HCN1-4* genes,^1^ that mediate the so-called “funny” current (I*_f_*) in the heart and “hyperpolarization-activated” current (I*_h_*) in the brain.

Even minor alterations in these channels’ voltage dependence and/or conductance have an adverse impact on membrane excitability leading to severe neuronal and/or cardiac dysfunction. Both gain- and loss- of HCN channel activity have been linked to several diseases, such as cardiac arrhythmias for HCN4, neuropathic pain, neurodevelopmental delay and epilepsy for HCN2 and neurodevelopmental delay and epilepsy for HCN1 channels.^2–5^

Over the past decade, both *de novo* and familial pathogenic variants have been identified in the *HCN1* gene of individuals presenting with a broad spectrum of epileptic disorders ranging from developmental and epileptic encephalopathy (DEE)^6^ to milder generalized epilepsies.^4,7,8^ Despite this growing recognition in clinical practice, clinicians still lack a functional framework to predict epilepsy risk, disease severity, and rational therapeutic direction based on the underlying channel defect. Even though a functional analysis of channel activity *in vitro* is available for a subset of variants, and preliminary classification efforts have been proposed,^7,9^ a comprehensive and systematic characterization of genotype-phenotype correlations in HCN1 is still lacking.

Most variants analyzed thus far were shown to yield HCN1 channels with a right (depolarizing) shift in the half-maximal activation voltage (V_1/2_), leading to facilitated channel opening. In many cases this was accompanied by an outright loss in the ability of the channel to fully close at rest, resulting in the emergence of a voltage-independent, “leak” current component, which we refer to hereon as “instantaneous”. These features have led to the prevailing interpretation that HCN1-related epilepsies predominantly represent gain-of-function mechanisms.^4,10^ One of the most outstanding questions is whether the opposite phenomenon, namely loss-of-function *HCN1* variants, lead to distinct clinical phenotypes.^4^ Another open question is whether the reduction in current density which is observed in mixed phenotypes along with facilitated channel opening, plays any role in determining the clinical outcome. Considering the complexity of HCN channel function, it is so far unknown whether reduced current density counteracts the gain-of-function imparted by a positive shift in the voltage response or if the mechanisms of voltage dependent gating determine genotype-phenotype correlations.

In this study, we provide a systematic and comprehensive classification of 43 *HCN1* variants, 15 of which were found in newly identified patients and are presented here for the first time. We further integrate this framework with detailed phenotypic data from an international patient cohort. By correlating the direction of channel dysfunction with epilepsy risk and severity, we propose a clinically meaningful model that may inform prognosis and guide precision therapeutic strategies.

We further explore conceptual approaches to therapy by testing the effects of newly available modulators of HCN channel gating. We show here that the most common defects, namely left and right shifts in the voltage dependence of the channel, can be corrected *in vitro* using allosteric compounds with the ability to modulate the interaction between the HCN1 voltage sensor domain (VSD) and the channel pore.

## Materials and methods

### Constructs and variants

Full-length human HCN1 (hHCN1) cDNA constructs, including eGFP- or TagRFP- mutants, were generated and expressed in HEK293T or HEK293F cells for functional studies. Mutations were introduced by site-directed mutagenesis using the QuickChange XL-II kit (Agilent Technologies) and sequences were verified. The mutations represent 15 variants found in newly identified patients, 1 novel variant found in an unborn fetus with finger abnormalities (E246K), as well as 5 variants present in the Clinvar database (W175R, N179Y, F186L, N200S, I206V). Nine additional variants were found in previously published patients but either lacked functional characterization (S100A, T172P, K261E, M379R, Y411C) or were previously tested using different heterologous expression systems (L157V, A387S, I380F, R590Q). Detailed construct design, cell culture conditions, transfection procedures and patient information are described in the Supplementary Material.

### Electrophysiology

Whole-cell patch-clamp recordings were performed on HEK293T or HEK293F cells as detailed in the Supplementary Material. Currents were recorded at room temperature using an ePatch (Elements srl) or a Dagan 3900A amplifier (Dagan Corporation); the latter signals were digitized using a Digidata 1550B (Molecular Devices). A detailed description of current recordings and analysis is provided in the Supplementary Material.

### Confocal microscopy

Confocal imaging was carried out on HEK293F cells, cultured and transfected as detailed in the Supplementary Material, using a Nikon Eclipse-Ti inverted confocal microscope. Samples plated on 35 mm glass Petri dishes were observed with a 60x 1.4 NA oil immersion objective (Nikon System). Details on image acquisition and data analysis are provided in the Supplementary Material.

### Analysis of clinical phenotypes

Probands carrying 49 novel or published *HCN1* variants were identified through an international multicentre collaboration. Both *de novo* and familial cases were included. Variants were annotated using NM_021072.4 and HGVS nomenclature. Standardized clinical data were collected, including epilepsy status, age at onset, syndrome, severity, neurodevelopmental outcome, and MRI findings. Epilepsy severity was classified as mild, moderate, or severe. For genotype-phenotype analyses, variants were divided into two groups based on functional characterization: loss-of-function (LOF) or non-LOF. The latter included gain-of-function (GOF), mixed effects (LOF/GOF) and variants with wild-type-like phenotype in heteromeric channel configuration. Associations were tested using logistic regression and contingency-table analyses. Detailed phenotypic definitions, functional classification, and statistical methods are provided in the Supplementary Material.

## Results

### Classification of *HCN1* variants into phenotypic classes

To avoid inconsistent results previously reported for different *in vitro* expression systems^4,6,9^ we expressed all variants in the mammalian cell line HEK293. In total, we collected functional data from 43 variants, of which 30 newly characterized here and 13 that had been previously characterized by us using the same experimental conditions.^4,9^ The phenotypic evaluation was done on heteromeric channels that mimic the heterozygous condition of all *HCN1* patients. For completeness, we also report the functional characterization of homomeric mutant channels.

We assigned the functional phenotypes to four classes, as illustrated in Fig. 1A:

**Figure 1.**
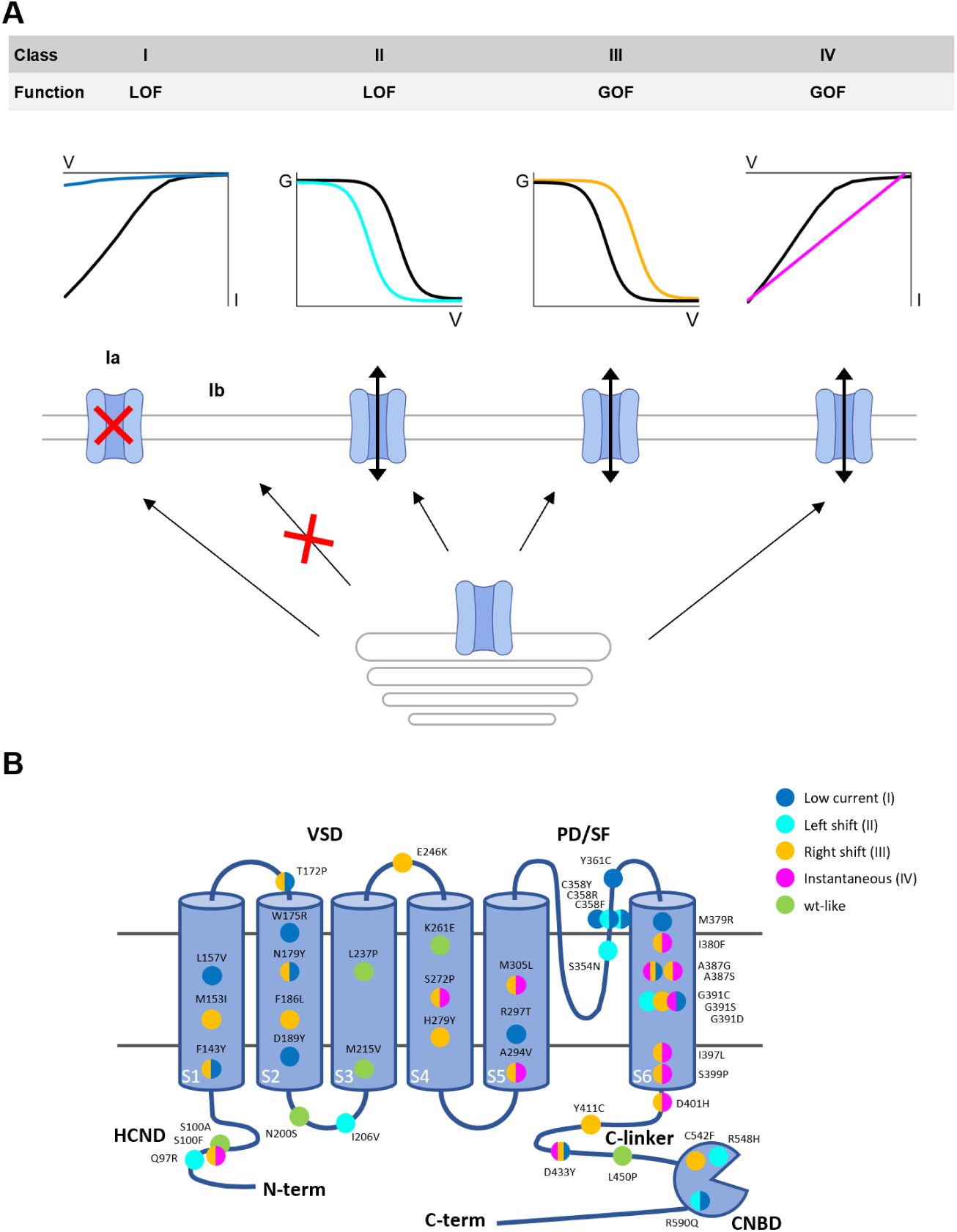
Functional classification of HCN1 variants. (**A**) Classes (I-IV) adopted to classify the phenotypic defects of heteromeric HCN1 mutant channels expressed in HEK293 cells and their subdivision by function: loss- (LOF) and gain-of-function (GOF). Middle: schematic representation of the functional defect in each class, shown as current/voltage (I/V) and conductance/voltage (G/V) relationships. Wild-type trace shown in black, mutant HCN1 shown in color. From left to right: I) low current density due to altered channel conductance (subclass Ia) or trafficking defects (subclass Ib); II) left shift of the voltage activation curve; III) right shift of the voltage activation curve; IV) presence of a voltage-independent (instantaneous) current component. Bottom: HCN1 channel expressed at the plasma membrane (double grey line) with ovals representing the intracellular secretory pathway. (**B**) Cartoon representation of one HCN1 monomer. Structural domains include: the cytosolic HCN domain at the N-term (HCND); six transmembrane domains (S1-S6), including the Voltage Sensor Domain (VSD, S1-S4) and the Pore Domain (PD, S5-S6) with the Selectivity Filter (SF); the cytosolic C-linker and the Cyclic Nucleotide Binding Domain (CNBD). All HCN1 variants functionally characterized in heterozygosis in HEK293 cells are shown as colored dots. Each mutation is color-coded as per legend: dark blue for low current density; light blue for left shift; orange for right shift; magenta for instantaneous current. Additionally, wild type-like variants (no functional alteration) are shown in green.

-class I, low or no current due to reduced conductance (sub-class Ia) or trafficking defects (sub-class Ib);

-class II, impaired gating: left shift in the activation curve;

-class III, impaired gating: right shift in the activation curve;

-class IV, presence of instantaneous (voltage-independent) current component.

We consider variants in class I and II as loss-of-function (LOF), as they show reduced current at physiologically relevant voltages. Variants in class III and IV are consequently interpreted as gain-of-function (GOF).

Fig. 1B shows all 43 variants, color-coded based on the classification system established above (classes I-IV); variants which exhibit a wild-type (wt)-like phenotype are shown in green. In several cases we could attribute a variant to a single class, such as W175R (I), I206V (II), H279Y (III) and many others. However, it was not possible to attribute univocally a variant to class IV (instantaneous current, magenta); this phenotype never manifested alone but was always associated with right shift and/or low current. Mutations showing “across-class” mixed phenotypes were also found in other combinations, although less frequently.

A few observations are immediately obvious: i) variants with significant effects on HCN1 channel function are found in almost all structural protein domains, including the extracellular and the intracellular loops connecting the transmembrane (TM) helices; ii) GOF mutations of class III (yellow) are found in all domains, with the notable exception of the selectivity filter (SF), while GOF mutations of class IV (magenta) are always mixed with a class III phenotype (with one exception) and are mostly found in the S5-S6 pore domain (PD). LOF mutations of class I and II (dark and light blue) are also widespread across domains with a prominent cluster in the SF; transmembrane helices S1 and S2 only carry variants with class I and class III phenotypes, iii) about half of the total mutations (18/43) are of mixed classes (more than one color).

Finally, what is particularly striking is the absence of variants in the S4-S5 linker in our collection of patients. This 12 aa linker (residues 281-293) forms the interfacial S4 helix that is created upon hyperpolarization during HCN channel gating.^11^ Remarkably, variants were found in positions immediately preceding (H279Y) and following (A294V) this segment, but none within the loop itself, suggesting a critical role in gating that likely makes this linker intolerant to variation.

### Detailed analysis of the functional properties of *HCN1* variants across classes

Fig. 2 shows the functional characterization of a representative member of each class and Fig. 3 shows the corresponding membrane localization analysis for Class Ia and Ib mutants. All other variants characterized in this study are presented in Supplementary Fig. 1, showing patch clamp and Supplementary Fig. 2 and 3, showing localization analysis. For each mutant, we provide V_1/2_, slope factors and steady-state current density values in Supplementary Tables 1 and 2, along with statistical analysis. Co-localization parameters are reported in Supplementary Tables 3 and 4.

**Figure 2.**
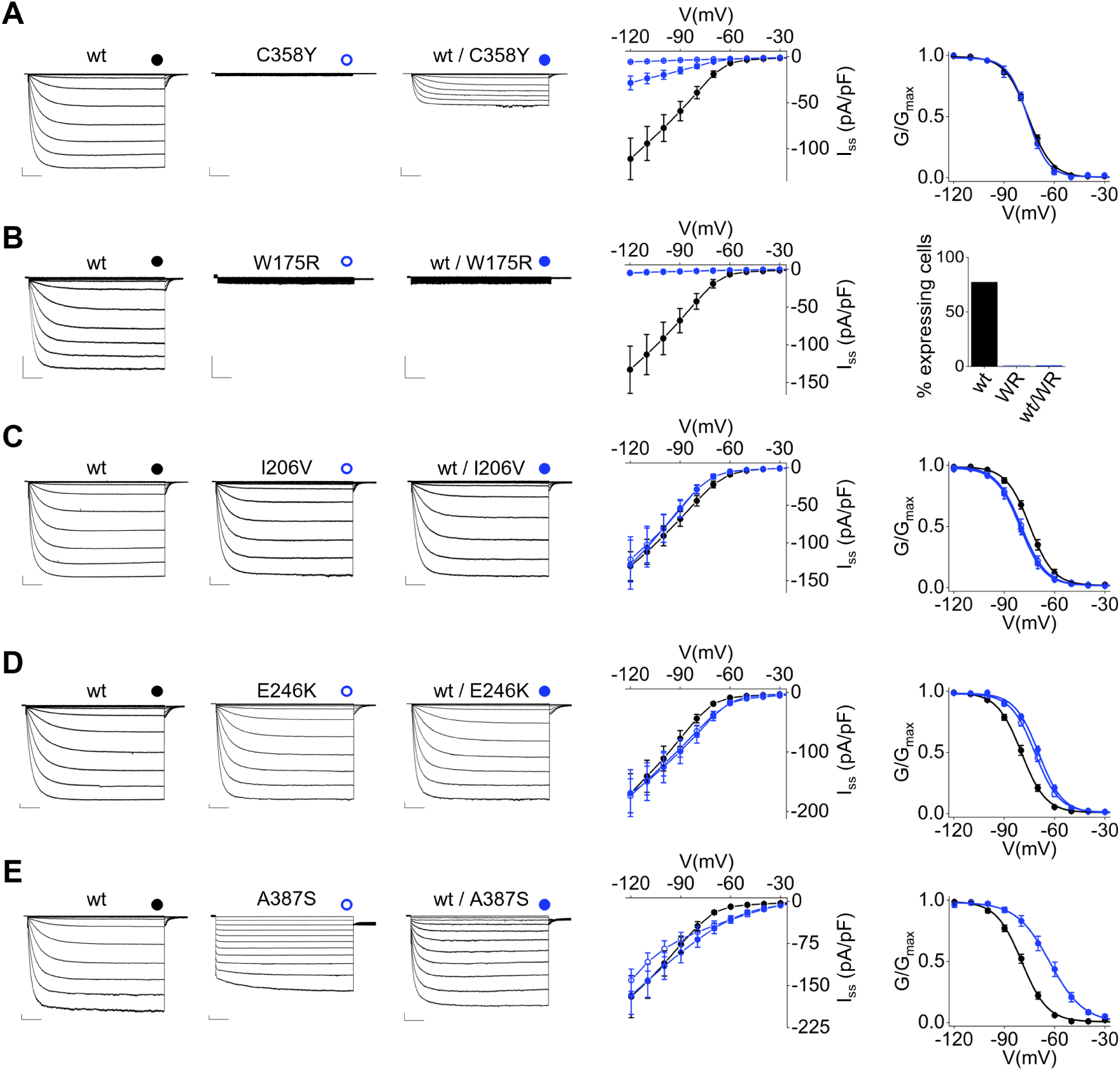
Functional properties of HCN1 variants representative of each class. Variants for class: Ia, C358Y (**A**); Ib, W175R (**B**); II, I206V (**C**); III, E246K (**D**); IV, A387S (**E**). Left: whole-cell currents recorded from HCN1 wild type channels (wt, black dot), homotetrameric (empty blue dot) and heterotetrameric mutant channels (solid blue dot). Traces shown from -20 mV to -120 mV. Scale bars: 250 pA and 500 ms. Middle: corresponding mean I/V relationships of steady-state current density (I_ss_, pA/pF). Data are mean ± SEM. Right: mean activation curves obtained from wt, homomeric and heteromeric channels. Data points are mean ± SEM. Data fit to the Boltzmann are plotted as solid lines. For variant W175R the histogram shows the percentage of cells with measurable current: wild type (wt, 77%), homomeric (WR, 0%) and heteromeric channel (wt/WR, 0%). I_ss_ values measured at -120 mV, V_1/2_, ΔV_1/2_, inverse slope factors (k) and number of cells (n) for each experiment shown are reported in Supplementary Tables 1 and 2 along with the details on statistical analysis.

**Figure 3.**
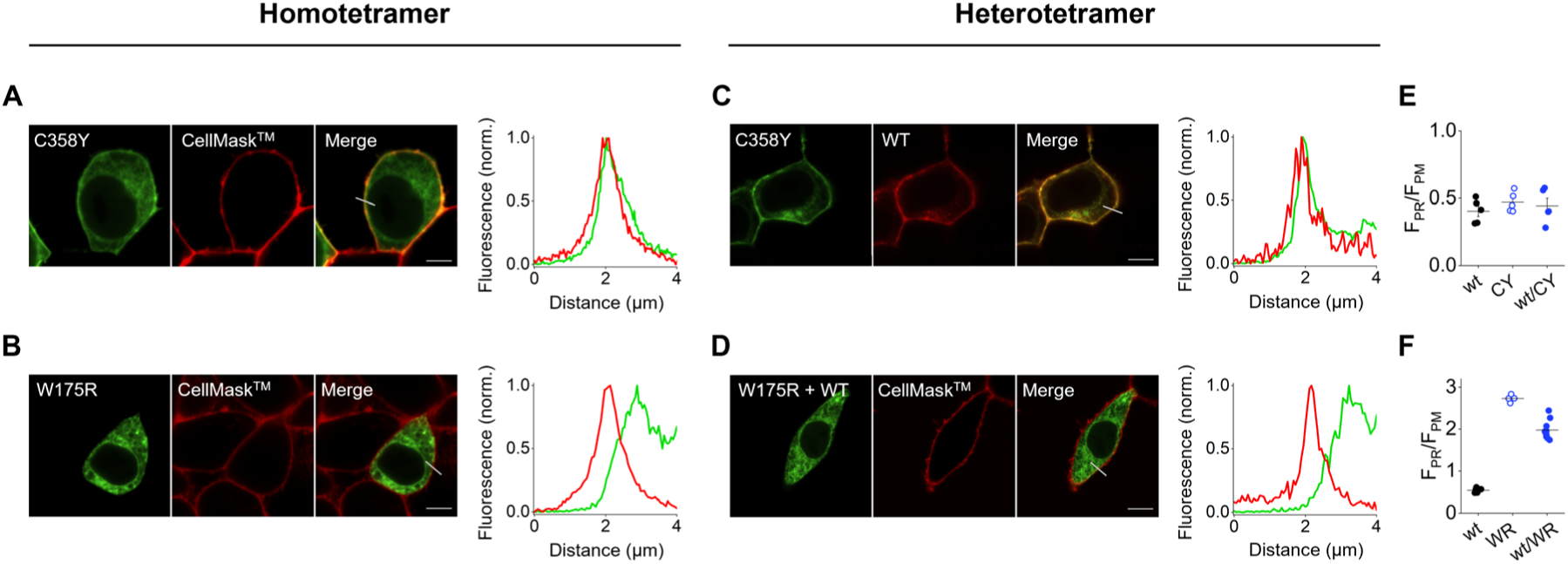
Confocal microscopy analysis of class I HCN1 variants. Confocal images of eGFP-HCN1 C358Y (Class Ia) and W175R (Class Ib) expressed alone (homotetramer) or co-expressed with Tag-RFP-HCN1 wild type chanels (heterotetramer) in HEK293F cells. Homotetramers, from left to right: eGFP-HCN1 C358Y (**A**) or W175R (**B**) (green), CellMask^TM^ plasma membrane dye (red) and merge of the two signals (yellow). Scale bar 5 µm. Right: representative normalized intensity profiles of green and red fluorescent signals encompassing the plasma membrane (grey line drawn on merged image). Overlapping red and green peaks indicate plasma membrane co-localization. Hetrotetramers, from left to right: **(C)** eGFP-HCN1 C358Y (green), Tag-RFP-HCN1 wt (red) and merge of the two signals (yellow). (**D**) eGFP-HCN1 wt + eGFP-HCN1 W175R (both green), CellMask^TM^ plasma membrane dye (red) and merge of the two signals (yellow). In this case, CellMask^TM^ was used to highlight the membrane profile, as the mutant prevents the wt channel from reaching the plasma membrane. Scale bar 5 µm. Right: representative normalized intensity profiles of green and red fluorescent signals along the grey line drawn on the merged image. Overlapping peaks indicate plasma membrane co-localization. (**E, F**) Mean fluorescent signal intensity expressed as the ratio of signal at the perinuclear ring (F_PR_) over that at the plasma membrane (F_PM_), F_PR_/F_PM_. Data shown as mean ± SEM. Quantitative analysis of F_PR_/F_PM_ ratio values, mean Pearson correlation coefficients (r), peak distance (PD) values and number of cells (n) are reported in Supplementary Tables 3 and 4 along with the details on statistical analysis.

Class 1, low or no current, is represented by two variants, C358Y (Class 1a), and W175R (Class 1b). C358Y does not conduct as a homotetramer (homo) but passes a small wt-like current as a heterotetramer (hetero, ∼ 25% of wt) (Fig. 2A, Supplementary Tables 1 and 2). Variant W175R yields no measurable hyperpolarization-activated current, neither alone nor when co-expressed with the wild type (Fig. 2B and Supplementary Tables 1 and 2).

Confocal imaging shows that variant C358Y reaches the plasma membrane (PM) where the homomeric channel co-localizes with the PM marker CellMask^TM^ (red) and the heteromeric channel with wild type subunits (green) (Fig. 3A,C). This is confirmed by the intensity profiles measured across the PM (grey line in Fig. 3) showing small peak distance (PD) value and a high Pearson correlation coefficient (r ∼ 1) (Supplementary Table 3). To quantify if mutant channels are retained intracellularly more than the wt, we calculated the ratio between the signal at the perinuclear ring (PR), a proxy for the endoplasmic reticulum, over that at the plasma membrane (F_PR_/F_PM_). The similar F_PR_/F_PM_ values obtained for mutant homo, hetero and wt channels, (homo = 0.5 ± 0.03, hetero = 0.4 ± 0.05, wt= 0.4 ± 0.03) indicate no specific retention of the mutant protein at the ER level (Fig. 3E and Supplementary Table 4). Thus, C358Y is properly synthesized and targeted to the membrane; still, it does not conduct current in homo and poorly conducts in hetero.

A different picture emerged for W175R. This variant does not reach the PM, neither as homo, nor as heteromeric channels (Fig. 3B,D). This is evident from the lack of co-localization with a PM dye, non-overlapping fluorescence intensity profiles and a low Pearson correlation coefficient (Fig. 3D,E and Supplementary Table 3). High values of the F_PR_/F_PM_ ratio (homo = 2.7 ± 0.03 and hetero = 2.0 ± 0.07) compared to the wt (0.5 ± 0.03) further indicate strong intracellular retention of the mutant protein, which leads to intracellular retention of wt subunits as well (wt/W175R) (Fig. 3F and Supplementary Tables 3 and 4). This analysis allowed to attribute wt/C358Y channels to subclass Ia (defect in conductance) and wt/W175R to subclass Ib (defect in trafficking).

To represent Class II (left shift) and Class III (right shift), we have chosen variants I206V and E246K, respectively (Fig. 2C,D). Heteromeric channels show the same amount of maximal current, evaluated at -120 mV, as the wt, but markedly shifted half activation voltages (V_1/2_). When expressed in hetero, wt/E246K channels are right shifted by +10.4 mV while wt/I206V are left shifted by -6.3 mV (Supplementary Tables 1 and 2). Thus, at physiologically relevant voltages (-90 to -40mV), I206V conducts less current (LOF) and E246K conducts more current (GOF) than wt channels.

Class IV defect, which is characterized by an instantaneous current component, was only found in association with other classes. Variant A387S (Fig. 2E) is representative of the most frequent finding (9 variants) where an instantaneous current occurs together with a right-shift of the activation curve (class III/IV). In one case (G391D) ^9^ the instantaneous current component was found together with a reduced current (Class Ia/IV) while in two others (A387G, D433Y) we found a 3-class defect comprising an instantaneous current, right shifted activation and low current (class Ia/III/IV) (Fig. 1B, Supplementary Fig. 1 and Supplementary Tables 1 and 2).

The instantaneous current component is evident in both the homomeric and heteromeric A387S channels (Fig. 2E). These channels conduct a small amount of current at voltages more positive than -60 mV, where the channel is typically closed, suggesting that the pore is slightly conductive at rest. In heterozygosis, a marked right shift in the activation curve (+16.5 ± 2.7 mV) is observed, which further increases the current compared to the wt, leading to a strong GOF phenotype (Supplementary Tables 1 and 2).

### Structural determinants of gain- and loss-of-function variants

Given the extensive understanding of structure-function correlates in HCN channels, it is of interest to examine structural explanations for the functional defects of mutants in relation to their position within the protein. This is especially relevant for variants located in the VSD, whose effects can be interpreted in the context of the well-characterized voltage-dependent gating mechanism of HCN1.^11–13^ Mutations F186L and D189Y on S2 and M153I on S1 affect residues of the “charge transfer center” (CTC) of the voltage sensor, a highly conserved structure formed by a hydrophobic cap and an hydrophilic binding site that controls the movement of positively charged amino acids in S4 across the membrane field.^14^ These positive charges are regularly spaced at every third amino acids along the S4 segment in nearly all members of the K_v_ family, and in HCN1 channels they are numbered as follows: R0^258^, K1^261^, S2^264^, R3^267^, R4^270^, R5^273^, R6^276^. As shown in Fig. 4A, M153 and F186 contribute to forming the “hydrophobic cap” to the binding site formed by D189 for the two arginine residues R3 and R4 that cross the CTC when they move downward in response to voltage. Mutations M153I and F186L reduce the hydrophobic side chains that form the CTC and facilitate voltage gating (right shift, class III). On the contrary, the loss of a negative charge due to mutation D189Y hampers gating (left shift, class II). Interestingly, charge inversion in K1, the second charged amino acid of the series, due to mutation K261E doesn’t display any detrimental phenotype in heterozygosis, while the homomeric channel does not reach the PM. This supports the notion that R3 and R4 only move through the transfer center during gating.^11^

**Figure 4.**
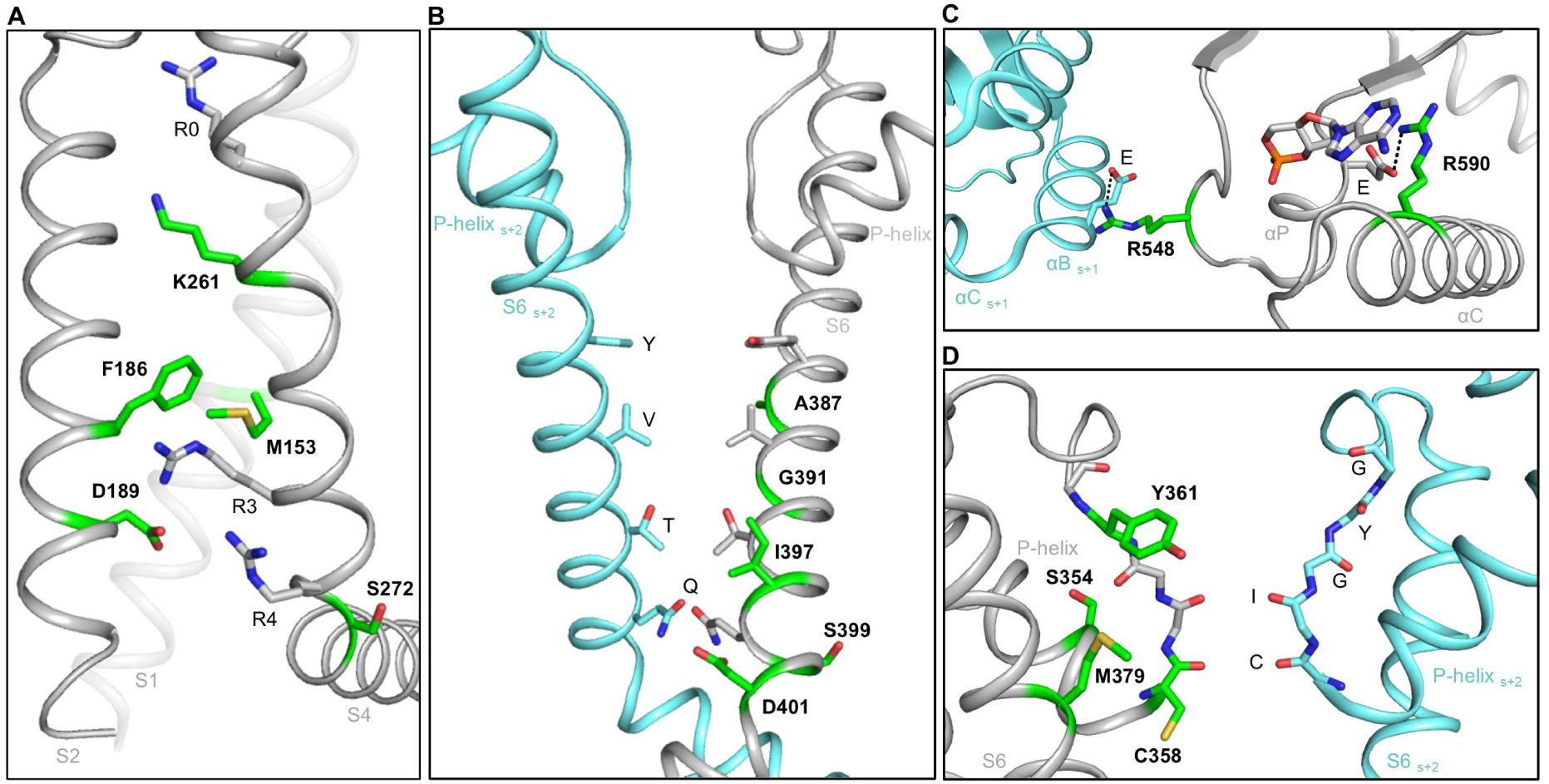
Structure-based molecular explanation of HCN1 dysfunction. Structure of the human HCN1 channel in a hyperpolarized conformation (PDB:6uqf), for clarity, only two of the four subunits are shown (grey and light blue). Residues found mutated in patients are shown in green. (**A**) Detailed view of the VSD showing the charge transfer center formed by residues F186, D189 on S2, and M153 on S1. Four of the charged residues on S4 (R0, K261, R3 and R4) are shown. Residue S272 marks the S4 breakpoint with the bottom half of S4 running parallel to the membrane plane. (**B**) Detailed view of the bundle crossing of the pore-lining S6 helices forming the hydrophobic gate YVTQ. (**C**) Detail of two arginine residues in the CNBD forming salt bridge interactions within the cAMP binding pocket (R590) and with the adjacent subunit (R548). (**D**) View of the selectivity filter showing the signature sequence CIGYG. Residues S354 and M379 in the pore helix that interact with the selectivity filter are also shown.

During gating, S4 helix kinks at serine 272 promoting e pore opening.^11^ Substitution with a proline in this position acts as a helix breaker, promoting pore opening at rest. This is reflected in the phenotype of the S272P variant which shows both a right shift in activation and an instantaneous current component^9^ (class III and IV) (Fig. 4A).

At the cytosolic end of S6, isoleucine 397 contributes to the hydrophobic contacts that keep the pore closed at rest (bundle crossing) (Fig. 4B). Substitution with the smaller leucine residue destabilizes the bundle, leading to facilitated gating (right shift, class III) and partly open pore at rest (instantaneous current, class IV). Similar effects were observed in other mutations of S6, I380F, A387G/S (Supplementary Fig. 1 and Fig. 2E), G391S/D,^4^ and of the C-linker, namely S399P and D401H,^9^ suggesting that these residues contribute to keeping the pore closed at rest. However, while D401H promotes pore opening, R297T does not, suggesting that in HCN1, these two residues, do not form a salt bridge, as previously proposed for HCN2.^15,16^

Finally, the LOF phenotype of R590Q (Supplementary Fig. 1) in the cyclic nucleotide binding domain (CNBD) can be explained by a decrease in cAMP affinity as this arginine interacts with the ligand^17^ (Fig. 4C). On the other hand, R548H disrupts an essential salt bridge with the adjacent CNBD that controls cooperativity of cAMP binding^18^ (Fig. 4C).

### Patients with *HCN1* variants but no epilepsy have loss-of-function variants

An unexpected finding of this study was that the largest group of mutations leading to LOF phenotypes clusters in the selectivity filter (C358F, C358R, C358Y, Y361C) or interacts with it from the pore helix (S354N) or S6 (M379R) (Fig. 5D). Cysteine 358 and tyrosine 361 are part of the so-called “signature sequence” of HCN1 (^358^CIGYG^362^). This sequence is highly conserved among HCNs as it forms the ion binding sites and controls selectivity and conductance.^19^ Residues S354 and M379 establish interactions with the SF that may stabilize it in the correct conformation. A first interesting finding is that mutations in this cluster lead to a left shift in the voltage dependence (Fig. 1). We interpret the effect of these mutations as evidence for non-canonical communication between the SF to the VSD.^20^

**Figure 5.**
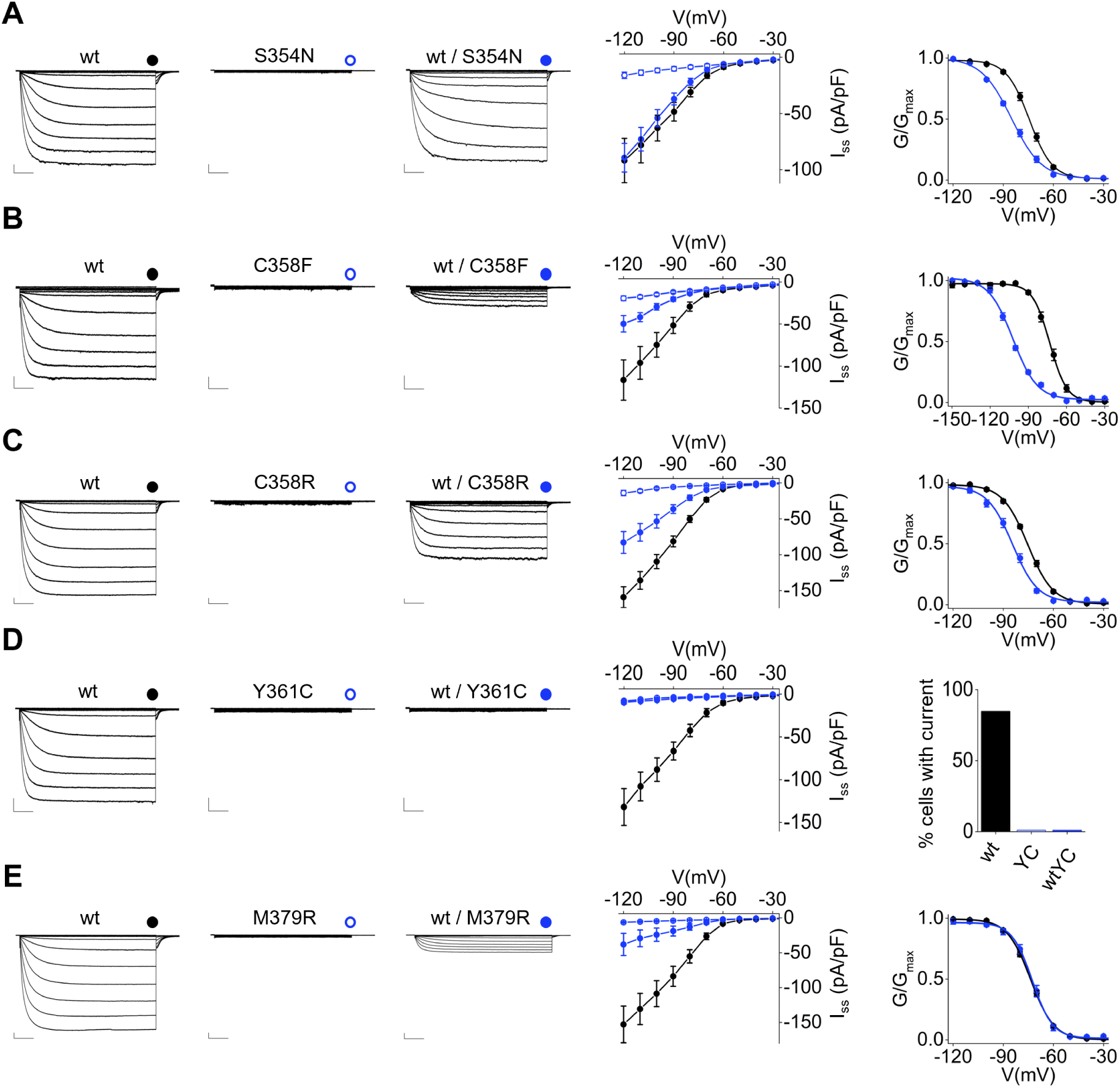
Functional properties of non-epileptic HCN1 pathogenic variants. Left: representative whole-cell currents of HCN1 mutants recorded in HEK293 cells: S354N (**A**), C358F (**B**), C358R (**C**), Y361C (**D**) and M379R (**E**). Cells were transfected with HCN1 wild type alone (wt, black dot), HCN1 mutant alone (homotetramers, empty blue dot) or both (heterotetramers, solid blue dot). Traces shown from -20 mV to -120 mV. Scale bars: 250 pA and 500 ms. Middle: corresponding mean I/V relationships of steady-state current density (I_ss_, pA/pF). Data are mean ± SEM. Right: mean activation curves obtained from HCN1 wt (black dots) and wt/mutant (solid blue dots). Data fit to the Boltzmann equation are plotted as solid lines. Data points are mean ± SEM. Where no activation curve could be obtained, the histogram with the percentage of cells expressing current is shown: (**D**) HCN1 wt (85%), Y361C (YC, 0%) and wt/Y361C (wt/YC, 0%). I_ss_ values measured at -120 mV, V_1/2_, ΔV_1/2_, inverse slope factors (k) and number of cells (n) for each experiment shown are reported in Supplementary Tables 1 and 2 along with the details on statistical analysis.

What makes this cluster even more interesting is that all six mutations were identified in patients with neurodevelopmental disorders (ND) but no evidence of epilepsy (see below and Supplementary Table 9). Because of this unexpected finding, we examined the functional properties of this set of mutants in Fig. 5; C358Y was previously shown in Fig. 2A. When expressed as homotetramers, none of the six mutants generated measurable currents. Trafficking defects were not observed, except for homotetrameric M379R (Fig. 3 and Supplementary Fig. 3). In contrast, when tested in combination with the wild type, heteromeric channels produced less current due to a negative shift in V_1/2_ (S354N, C358F, C358R), a decrease in maximal conductance (C358Y, Y361C, M379R) or a combination of both. Overall, these functional properties are not strikingly different from other LOF variants which have been found in patients presenting with epilepsy, such as class 1a wt/R297T,^9^ class II R548H (Supplementary Fig.1) and class Ia/II R590Q (Supplementary Fig.1). The fact that LOF mutations which are not linked to epilepsy cluster in the pore/filter region suggests a possible compensation mechanism arising from cellular context that was not analysed here.

### The Therapeutic Potential of Nanobodies and Peptides

After classifying *HCN1* variants into different functional groups, we searched for treatments which could compensate for specific functional defects. Aiming to develop allosteric modulators of HCN channels that would affect voltage dependent gating without reducing the maximal channel conductance, we tested the functional effects of nanobody 6 (NB6), a biologic we previously isolated from a synthetic library as an activator of HCN4.^21^ NB6 also recognizes HCN1 but not HCN2 or other channels including hERG. Incubation with 20 µM NB6 induces an ∼8 mV depolarizing shift in the voltage dependent activation of HCN1, effectively increasing the current at non-saturating voltages (Fig. 6A and Supplementary Tables 5 and 6). Thus, NB6 represents the first known HCN1 channel activator and therefore deemed suitable for the rescue of *HCN1* LOF variants of class II. We tested NB6 on the R548H variant which is left shifted by ∼5 mV when expressed in heterozygosis. NB6 indeed reverted the effects of the mutation, restoring wt-like values of voltage dependence and kinetics (Fig. 6B and Supplementary Tables 5 and 6). Similarly, NB6 rescued the LOF phenotype of the non-epileptic variant wt/S354N (Fig. 6C and Supplementary Tables 5 and 6). Notably, NB6 exerts its effect by binding to the extracellular side of the channel, while mutations R548H and S354N are located in an intracellular domain and the pore, respectively.

**Figure 6.**
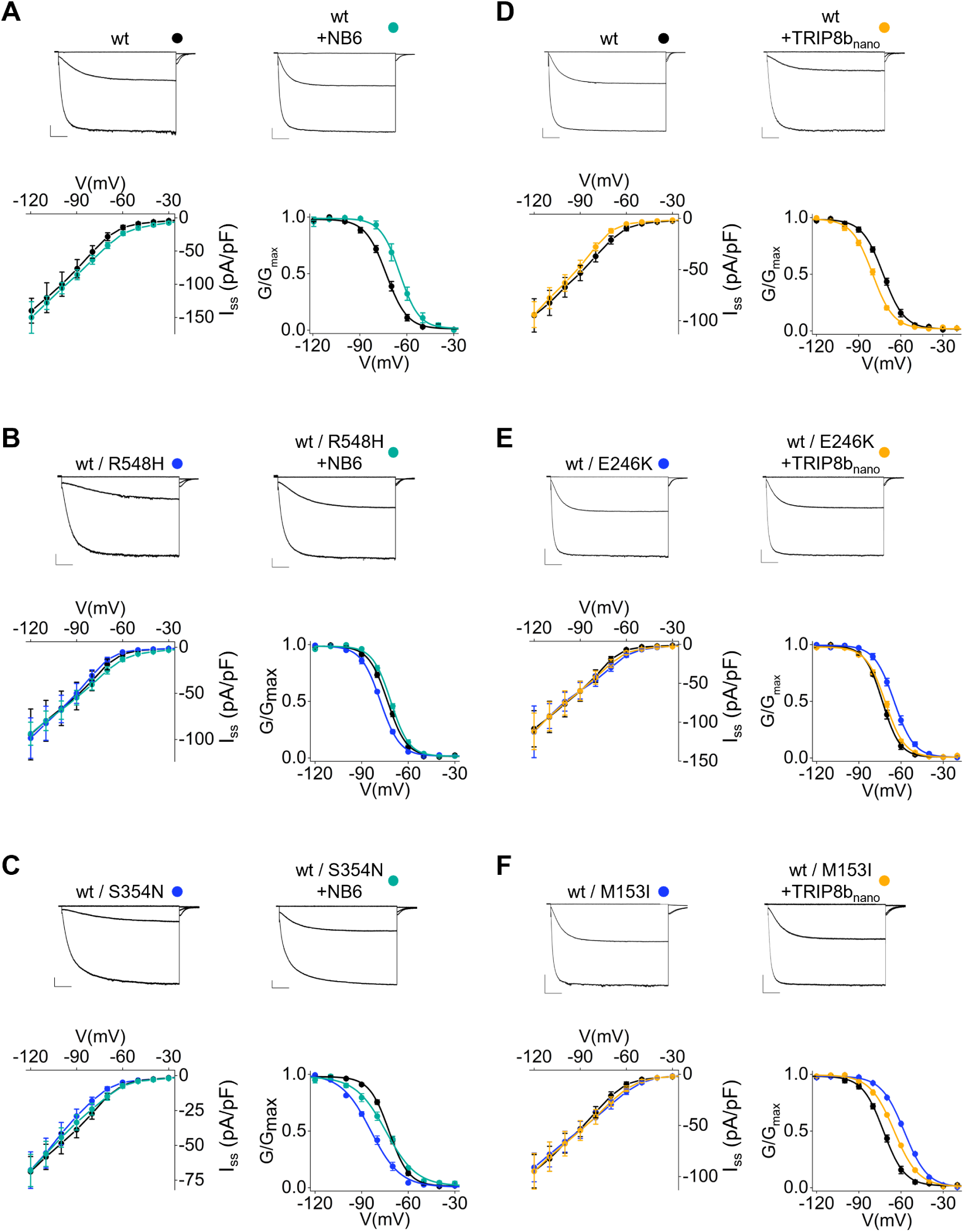
Peptide-based rescue of class II and III HCN1 pathogenic variants. Functional testing of HCN1 wt (black dot) and HCN1 wt/mutant (blue dot) in control solution or treated with 20 µM NB6 in the extracellular solution (+NB6, teal): HCN1 wt (**A**), wt/R548H (**B**) and wt/S354N (**C**); or 10 µM TRIP8b_nano_ (+TRIP8b_nano_) in the pipette solution: HCN1 wt (**D**), wt/E246K (**E**) and wt/M153I (**F**). For each panel, top: representative whole-cell current traces shown for -20, -80 and -120 mV voltage steps. Scale bars: 250 pA and 500 ms. Bottom left: mean I/V relationships of steady-state current density (I_ss_, pA/pF). Data are mean ± SEM. Bottom right: mean activation curves, with data fit to the Boltzmann equation plotted as solid lines. Data points are mean ± SEM. I_ss_ values measured at -120 mV, V_1/2_, ΔV_1/2_, inverse slope factors (k) and number of cells (n) for each experiment shown are reported in Supplementary Tables 5 and 6 along with the details on statistical analysis.

As a second approach to allosteric modulation of HCN1 with peptide binders, we employed TRIP8b_nano_, a rationally designed peptide (40 aa) that binds to the CNBD. TRIP8b_nano_ prevents cAMP binding and thereby induces a hyperpolarizing shift in channel activation.^22^ When added in the patch pipette, TRIP8b_nano_ causes an ∼8 mV hyperpolarizing shift in the activation curve of wild type HCN1, effectively counteracting the effect of endogenous cAMP in HEK293 cells (Fig. 6D and Supplementary Tables 5 and 6). When tested on variant E246K, a GOF variant of class III, 10 µM TRIP8b_nano_ fully rescued the 8 mV depolarizing shift caused by the mutation expressed in heterozygosis (Fig. 6E and Supplementary Tables 5 and 6). Similarly, TRIP8b_nano_ was able to partially correct the altered voltage dependence of variant wt/M153I, another GOF variant which exhibits a markedly right shifted (+17 mV) activation (Fig. 6F and Supplementary Tables 5 and 6).

These proof-of-concept experiments underscore the power of exploiting defined functional classes of *HCN1* variants to guide the rational selection of modulators suited to rescue the specific alterations observed in each group.

### Small-molecule allosteric inhibitors of HCN channels partially restore normal gating properties in HCN1 channels with gain-of-function mutations

Among epilepsy-associated *HCN1* variants, we observed a high prevalence of class III phenotypes with a right shift in the activation curve of the channel that facilitates channel opening in response to hyperpolarizing voltage changes. This impact on gating, which is often accompanied by a voltage-independent “leak” current, prompted us to ask whether known HCN channel inhibitors may help curb the effects of such gain-of-function variants. Currently, there is only one drug that targets HCN channels and is available for clinical use, namely ivabradine (sold as Corlanor in the United States and as Procoralan in Europe). When tested on mutation A387S, a GOF variant characterized by a strong instantaneous component, ivabradine (30 µM) was as effective as the known HCN blocker Caesium (5mM) (Supplementary Fig. 4A-C) in reducing both instantaneous and time dependent current components to almost zero. Ivabradine is used safely and effectively for the treatment of chronic stable angina pectoris.^23^ However, its limited ability to cross the blood-brain barrier and its potential bradycardic side-effects prevent its use for the treatment of neurological and neuropsychiatric conditions.^24,25^ Thus, there has been an ongoing effort to identify new compounds that can target HCN channels in the brain.

One such compound, originally developed by Organon for the treatment of depression and sleep disorders (WO1997040027A1, US20110092486A1) is Org-34167. Previous studies using heterologous expression of wild type and mutant HCN1 subunits in *Xenopus* oocytes have shown that Org-34167 inhibits HCN channels through an allosteric mechanism of action, modulating the interaction between the voltage sensor and the pore domain^26^; this action reduces the instantaneous “leak” current caused by several DEE-associated variants.^27^ Since the plasma membrane composition in *Xenopus* oocytes is considerably different from that of mammalian cells^28^ and HCN channels are extremely sensitive to lipid environment,^29^ we first sought to confirm these effects in HEK293 cells. Testing of 60 µM Org-34167 on homomeric A387S channels confirmed its ability to markedly reduce the instantaneous current component without completely blocking the slow activating voltage-dependent component (Supplementary Fig. 4D-F). However, control experiments revealed significant off-target effects of this compound. First, we observed a prominent reduction in the amplitude of endogenous outward currents, presumably carried by K^+^, when Org-34167 was tested in untransfected HEK293F cells (Supplementary Fig. 5A). Second, Org-34167 showed a profound effect on cardiac hERG channel activity, with a ten-fold decrease in steady state current compared to vehicle-treated cells (Supplementary Fig. 5B). Finally, Org-34167 was active on Na_V_1.5 sodium channels (Supplementary Fig. 5C) causing an almost complete disappearance of steady-state current. These data suggest widespread effects of this drug on other ion conductances, which critically compromise the use of Org-34167 as a selective therapeutic tool for HCN-associated disorders.

Another series of compounds targeting HCN channels with the ability to cross the blood-brain barrier was developed by Johnson & Johnson.^30^ Compound J&J12e in this series displayed nanomolar potency on HCN1 when tested by automated patch-clamp in HEK293 cells (IC_50_ = 58 nM).^31^ When we tested J&J12e in untransfected HEK293F cells, using 50 nM, a concentration near its IC_50_ for HCN1 channels, we did not observe any effect on endogenous outward currents (Supplementary Fig. 5D). No effects were noted when we tested this compound on hERG channels or Na_V_1.5 channels either (Supplementary Fig. 5E,F), suggesting a much higher specificity of J&J12e for HCN channels, compared to Org-34167.

To characterize more in detail the actions of J&J12e, we carried out a series of experiments testing a range of drug concentrations on wt HCN1 channels. As shown in Fig. 7, increasing the concentration of J&J12e from 1 nM to 1 mM progressively reduced HCN1 steady-state current (Fig. 7A,B and Supplementary Table 7). Activation curves obtained at concentrations equal or higher than 10 nM showed a gradual, progressively larger shift of V_1/2_ towards more negative values as drug concentration was increased (10 nM, ΔV_1/2_ = -5.0 ± 1.6 mV; 25 nM, ΔV_1/2_ = -12.1 ± 1.8 mV; 50 nM, ΔV_1/2_ = -15.2 ± 2.0 mV) (Fig. 7C and Supplementary Table 7). To determine whether J&J12e also affects HCN1 conductance, independent of its action on the channels’ voltage response, we next measured the drug’s effect on maximal tail current amplitude. As shown in Fig. 7D, the observed reduction in steady-state current is due to a decrease in channel conductance that occurs independently of J&J12e’s action on the channel’s voltage response (Supplementary Table 7).

**Figure 7.**
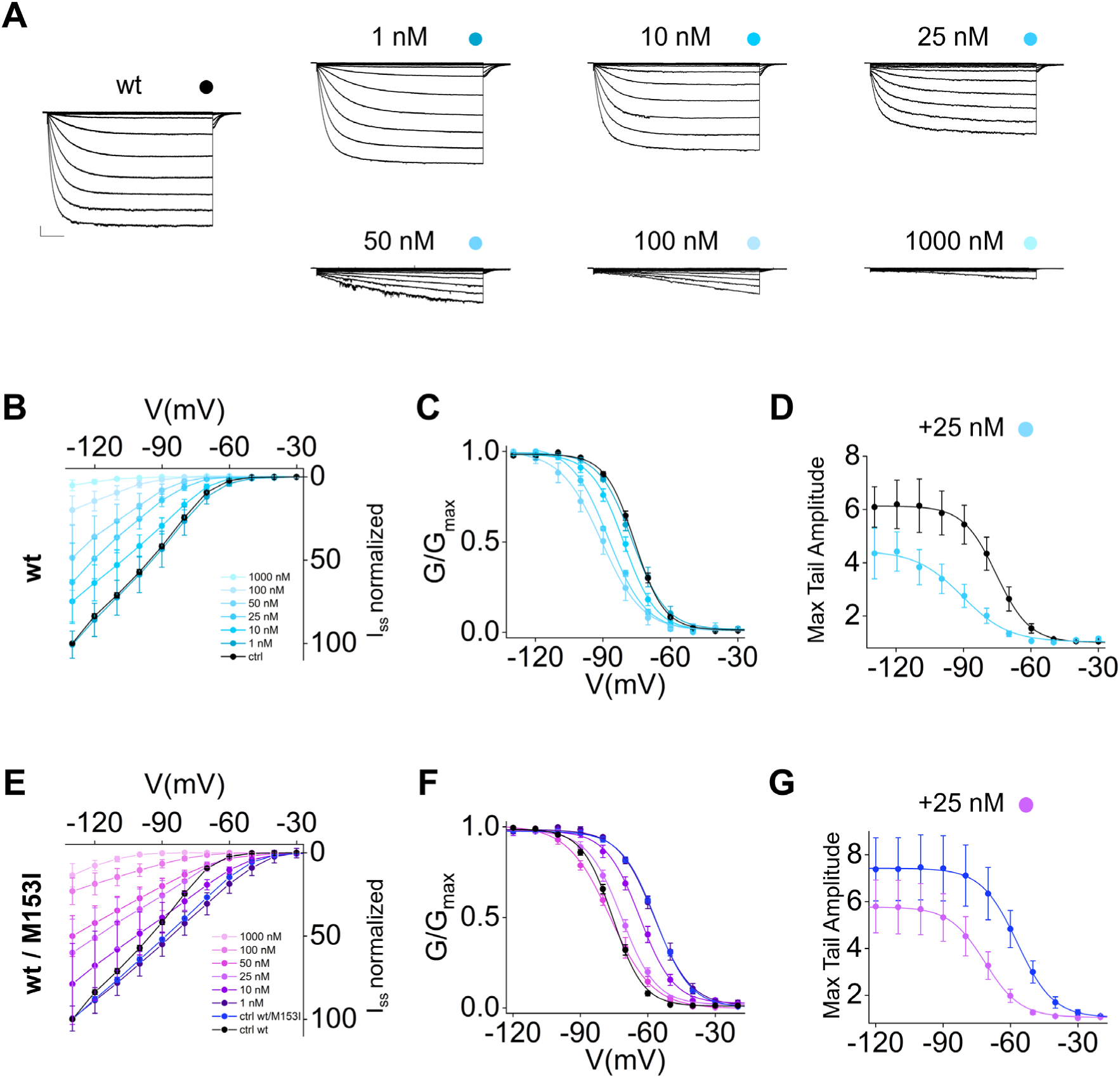
J&J12e rescues pathogenic variant M153I. (**A**) Representative whole-cell currents of HCN1 wild type treated with increasing concentrations of J&J12e. Traces shown from -20 mV to -120 mV. Scale bars: 250 pA and 500 ms. (**B**) Mean I/V relationship of HCN1 wild type treated with increasing concentrations of J&J12e. Data points represent current density values (in pA/pF) normalized to non-treated controls (I_ss_ normalized). Data are mean ± SEM. (**C**) Mean activation curves obtained from HCN1 wt in control solution (black dots), or in presence of J&J12e (light blue dots, color-coded as in legend). Data fit to the Boltzmann equation are plotted as solid lines. Data points are mean ± SEM. (**D**) Mean normalized maximal tail current amplitude obtained from HCN1 wt in control solution (black dots) or in the presence of 25 nM J&J12e (light blue dots). Data fit to the Boltzmann equation are plotted as solid lines. Data points are mean ± SEM. (**E**) Mean I/V relationship of HCN1 wt / M153I treated with increasing concentrations of J&J12e. Data points represent current density values (in pA/pF) normalized to non-treated controls (I_ss_ normalized). Data are mean ± SEM. (**F**) Mean activation curves obtained from HCN1 wt /M153I in control solution (blue dots), or in the presence of J&J12e (purple dots, color-coded as in legend). HCN1 wild type shown in black for reference. Data fit to the Boltzmann equation are plotted as solid lines. Data points are mean ± SEM. (**G**) Mean normalized maximal tail current amplitude obtained from HCN1 wt /M153I in control solution (blue dots) or in presence of 25 nM J&J12e (purple dots). Data fit to the Boltzmann equation are plotted as solid lines. Data points are mean ± SEM. I_ss_ values measured at -120 mV, V_1/2_, ΔV_1/2_, inverse slope factors (k), max tail amplitude values and number of cells (n) for each experiment shown are reported in Supplementary Table 7 along with the details on statistical analysis.

Overall, these experiments show that J&J12e can be used as an effective tool to fine tune the response to voltage of HCN1 channels in a concentration dependent manner. It has therefore the potential to correct the functional properties of mutants with depolarized V_1/2_, albeit at the expense of a somewhat reduced current density. Since our genotype-phenotype analysis suggests that decreased current density is not “per se” a critical determinant of epilepsy (Fig. 5), we surmise that this additional effect should not worsen the epileptic phenotype. On the other hand, channel inhibitors which act as pore blockers without altering the channels’ voltage response, such as ivabradine or the new series of HCN1-specific compounds recently developed by Roche,^31^ do not seem poised to address the more defining element of gain-of-function epilepsy-associated *HCN1* variants, which is their altered response to voltage.

Given these premises, we sought to test the effects of J&J12e on a prototypical *HCN1* variant which displays a very large positive shift in V_1/2_ but no instantaneous “leak” current component, namely M153I.^9^ Two independent patients carrying the M153I mutation *de novo* present with DEE along with mild ID and language delay.^4^ Consistent with prior studies, heteromeric HCN1 wt/M153I channels displayed a +18 mV shift in V_1/2_ (Fig. 7E-G and Supplementary Table 7). Titration of J&J12e demonstrated that concentrations between 25-50 nM were able to essentially restore normal voltage response in mutant channels, shifting the half-maximal activation voltage back to wild type levels (wt HCN1, V_1/2_ = -75.9 ± 0.8 mV; M153I/wt + 25 nM, V_1/2_ = -71.8 ± 1.2 mV; M153/wt + 50 nM, V_1/2_ = -76.6 ± 1.6 mV) (Fig. 7F and Supplementary Table 7), albeit with a progressive reduction in steady state current (Supplementary Table 7). As shown in Fig. 7G, measures of max tail current amplitude demonstrated a modest ∼22% reduction in current density at 25 nM drug concentration, which increased to ∼ 32% at 50 nM (Supplementary Table 7).

The combination of the two effects, left shift and max current reduction, induced by the drug, efficiently counteracts the GOF phenotype of wt/M153I channels. Close scrutiny of Fig. 7E further highlights that in the concentration range 10-50 nM, J&J12e restores wt-like current values of the mutant in the physiological-relevant voltage range between -90 mV and -60 mV.

In conclusion, while further studies will be needed to determine whether the stability and pharmacokinetic properties of J&J12e are appropriate for chronic use *in vivo*, the experiments above demonstrate that a class of compounds which can allosterically modulate the voltage response of HCN channels may provide a valuable precision medicine tool for the treatment of HCN1-linked epilepsies.

### Clinical characteristics of the cohort and genotype-function-phenotype correlations

#### Cohort overview

We evaluated 49 probands (29 female, 20 male) carrying 37 distinct *HCN1* variants; of these patients, 17 carried novel variants (Supplementary Table 9). Several recurrent amino-acid substitutions were identified, including M153I, S272P, M305L, G391D (each found in two unrelated patients) and C358R, I380F, A387S, G391S (each found in three unrelated patients). We also identified multiple different substitutions occurring at some sites, most notably C358F/R/Y in the selectivity filter and G391C/D/S in the S6 pore domain. Of the 49 probands, 42 patients (85.7%) presented with epilepsy whereas seven (14.3%) had no epilepsy (Fig. 8A and Supplementary Tables 8 and 9). Two probands were found to each carry two separate *HCN1* variants and were excluded from further analysis (Supp. Table 9, patient ID#10 and patient ID#12).

**Figure 8.**
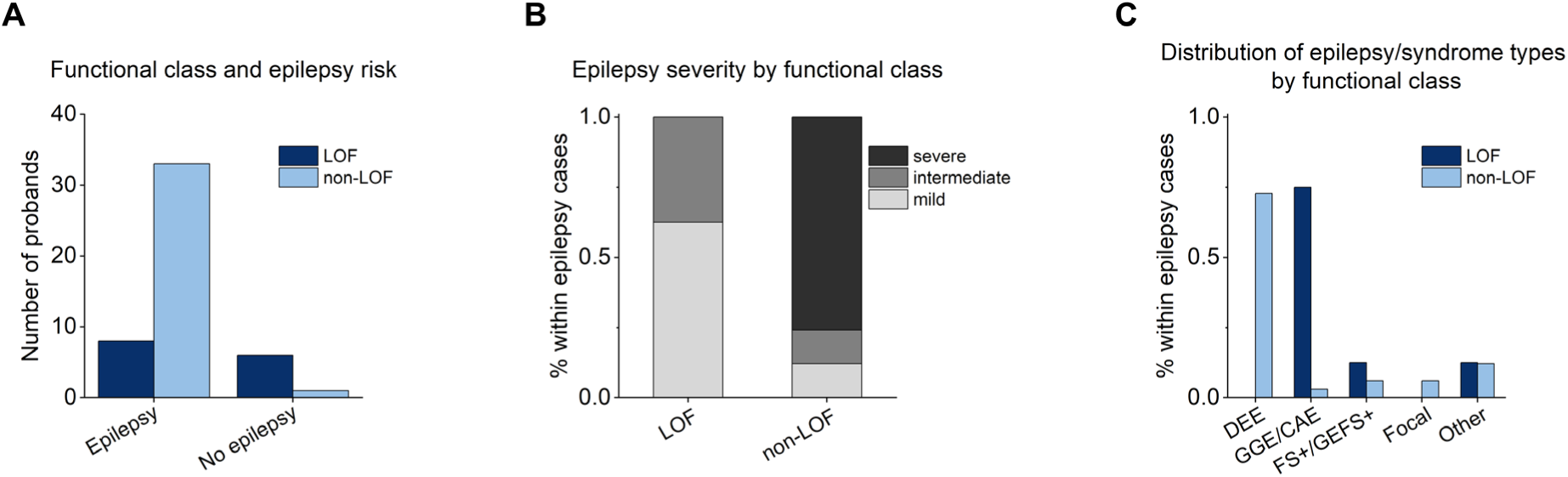
Functional class and epilepsy phenotype in HCN1-related epilepsy. (**A**) Epilepsy occurrence according to functional class. LOF variants were enriched among individuals without epilepsy, whereas non-LOF variants predominated among epilepsy cases (OR 0.04, 95% CI 0.004-0.38, p = 0.005). (**B**) Epilepsy severity among probands with epilepsy (n = 41). Severity distribution differed significantly between functional classes (Fisher-Freeman-Halton exact test, p = 4 × 10⁻⁵); severe phenotypes were confined to the non-LOF group. LOF (n = 8), non-LOF (n = 33). (**C**) ILAE epilepsy/syndrome distribution among probands with epilepsy. Functional class was significantly associated with epilepsy/syndrome category (Fisher-Freeman-Halton exact test, p = 2.9 × 10⁻⁵), with DEE restricted to the non-LOF group and generalized/absence-spectrum epilepsies enriched in LOF carriers. Details on patient distribution among classes and phenotypic severity are shown in Supplementary Tables 8 and 9.

#### Functional class and epilepsy risk

Among the 41 probands with epilepsy analyzed, eight (19.5%) carried LOF variants and 33 (80.5%) carried non-LOF variants, including 24 GOF, six associated with both LOF and GOF effects (LOF/GOF) and three with wt-like behavior when tested in our heterologous expression assay (Figure 1). Conversely, among the seven probands without epilepsy, six (85%) carried LOF variants and one (15%) carried a wt-like variant. A logistic regression model demonstrated a significant association between functional effect and epilepsy occurrence. LOF variants were associated with significantly lower odds of epilepsy compared to non-LOF variants (OR 0.04, 95% CI 0.004-0.38, p = 0.005) (Fig. 8A and Supplementary Tables 8 and 9).

#### Functional class and epilepsy severity

Severity distribution differed significantly according to functional class (Fisher-Freeman-Halton exact test, p = 4 × 10⁻⁵). No severe cases were observed among LOF variants (0/8), whereas severe phenotypes predominated in the non-LOF group (25/33, 75.8%). When dichotomized (severe vs non-severe), this association remained significant (Fisher’s exact test, p = 1.3 × 10⁻⁴), with a markedly reduced odds of severe epilepsy among LOF carriers (OR 0.02, 95% CI 0.001–0.38). Given the limited sample size and the presence of complete separation, these findings should be interpreted cautiously (Fig. 8B and Supplementary Tables 8 and 9).

#### Functional class and epilepsy phenotype (ILAE groups)

epilepsy types and syndromes were grouped according to ILAE classification into DEE, GGE/absence-spectrum, focal epilepsies, FS+/GEFS+, and other/unclassified categories. The distribution differed significantly according to functional effect (Fisher–Freeman–Halton exact test, p = 2.9 × 10⁻⁵). All DEE occurred in the non-LOF group (24/24), whereas none were observed among LOF variants. Conversely, GGE/absence-spectrum epilepsies were markedly enriched in the LOF group (6/8 LOF vs 1/33 non-LOF). No significant associations were observed for focal epilepsies, FS+/GEFS+, or other/unclassified categories (Fig. 8C and Supplementary Tables 8 and 9).

Given the limited sample size and the low frequency of several individual syndrome categories, these findings should be interpreted with caution and considered exploratory.

## Discussion

Pathogenic variants in *HCN1* are now an established cause of a spectrum of epilepsies, ranging from severe Developmental and Epileptic Encephalopathy (DEE) to milder forms like Genetic Generalized Epilepsy (GGE).^4,6,9,32,33^ The widespread inclusion of *HCN1* in diagnostic gene panels has led to a growing number of variant identifications in routine clinical practice. However, for many clinicians and families the detection of an *HCN1* variant still provides limited guidance regarding prognosis, epilepsy risk, and potential therapeutic options.

In channelopathies involving the cystic fibrosis transmembrane regulator (CFTR), the implementation of a classification system for pathogenic variants has greatly advanced the development of targeted, personalized therapies^34^ and enabled the prediction of disease severity for newly identified variants.^35^ Inspired by this framework we sought to develop a mechanistically informed model capable of linking direction of channel dysfunction to clinical phenotype in HCN1-related disorders. Based on a range of functional defects caused by a large number of *HCN1* variants, we defined four distinct classes encompassing defects in channel trafficking, conductance, and gating.

To ensure consistency across variants, wild-type and mutant channels were expressed in the same heterologous system, the mammalian HEK293 cell line. Functional parameters and cellular localization were examined using the same criteria and standardized protocols. This approach was essential to avoid functional disparities arising from differences in expression systems and analysis, as previously reported.^9^ To reflect the situation in patients, our functional classification was based on the properties of heteromeric channels, as all *HCN1* pathogenic variants identified in patients to date are dominant with probands heterozygous for the mutation.

The four-class system that we established does not merely describe biophysical diversity but provides a framework that can be directly integrated with patient-level clinical data. In the absence of any experimental evidence of mutations which corrupt early stages of HCN1 protein synthesis, our classification does not include a specific category for these defects.

A key observation is that approximately half of the variants show mixed phenotypes, i.e. combinatorial defects that span more than one functional class. The most prominent example of such mixed phenotypes is class IV, in which the prevalence of an instantaneous current is never found alone but instead occurs in combination with either a right shift in voltage dependence (class II) or a reduced current amplitude (class I).

Importantly, we also identified five DEE-associated variants that we classified as wt-like as they did not display detectable differences from wild type channels when expressed in heterozygosis. It is possible that these variants are benign, with the patient’s phenotype instead resulting from mutations in other genes. Of note 3/5 of these wild type-like variants (L237P, K261E, L450P) resulted in LOF when expressed as homomers, suggesting impaired subunit function. An alternative possibility is that their impact is sensitive to the balance of mutant vs wild-type HCN subunits present in native conditions, including potentially HCN2 subunits which are co-expressed with HCN1 in a variety of cortical neuron classes.^1^ Finally, our findings may reflect an inherent limitation of HEK293 cells as a test system for HCN1 channels. As with all heterologous expression systems, they lack typical neuronal features and will be opaque to important cellular functions (e.g. cell compartment trafficking and/or modulation by neuron-specific intracellular factors).

### Correlation between variant topology and functional aberrations

Scrutiny of the topology and the frequency of the mutations analyzed here shows no obvious pattern with respect to the four classes. The data nonetheless reveal some trends. It is well known that voltage-dependent opening of the HCN pore is allosterically controlled by membrane voltage and ligand binding to the CNBD.^36^ It is hence not surprising that GOF mutations belonging to Class III (right shift) were the most abundant (20/43) and were found in all protein domains, both transmembrane and cytosolic. Class I (no or low current) was the second most represented group (14/43), with two clear hotspots in the SF and in the S2 transmembrane domain. Class II (left shift) was relatively less abundant (8/43). These variants were not observed in the VSD but clustered within the SF or were found in the cytoplasmic domains that control cAMP binding (CNBD) and efficacy (HCND and S2-S3 loop).^37,38^ The latter finding is not surprising, given that cAMP binding shifts the channel’s voltage dependence to the right, an effect observable in HEK cells in basal conditions. ^37,38^

Present knowledge of structure-function correlations in HCN channels allows, in many cases, to explain the effects of several specific mutations in molecular terms. While many functional features of the analyzed mutants are consistent with known structure-function correlates in HCN channels, our analysis also led to the discovery of previously unknown interactions. Specifically, the observation that mutation M153I causes a similar functional phenotype to F186L allowed to identify the structural evidence that M153 contributes to the formation of a hydrophobic plug. This plug interacts with the gating arginines of S4 and controls their translocation during voltage sensing.^39,40^

A major clinically relevant finding of this study is the identification of a subgroup of loss-of-function variants clustered within the selectivity filter that were consistently associated with neurodevelopmental impairment in the absence of epilepsy. In our analysis we find that these heteromeric channels show reduced or no current (class Ia: C358Y, Y361C, M379R), left shift (class II: S354N) or mixed low current and left shift (class Ia/class II: C358F, C358R). These altered functional features are however similar to those found in variants associated with epilepsy, such as D189Y, R297T (class Ia); Q97R, I206V, R548H, G391C (class II); R590Q (class Ia/II). The only trait distinguishing these mutants from other LOF variants within the same classes is their location in the selectivity filter or at positions that interact with it. A reasonable hypothesis is that these mutations alter channel conductance and/or ion selectivity in ways that are not detectable under the standard experimental conditions adopted in this screening. However, this hypothesis does not account for the non-epileptic phenotype of the S354N variant which exhibits a left shift in activation without any reduction in maximal current. This observation underscores the role of the SF in controlling the voltage-dependent activation of HCN channels, an interesting functional feature that alone cannot explain why this cluster of mutations does not result in epilepsy.

### Classification of HCN1 mutant channels allows targeted correction of channel dysfunction

Here, we provide proof-of-principle evidence demonstrating that the aberrant functional properties of class II, III and IV variants can be corrected in a class-specific manner, highlighting the value of a mechanistically informed classification. The left shift in the activation curve of Class II variants wt/R548H and wt/S354N is corrected by NB6, even though these two mutations reside in domains that are structurally distant from the NB6 binding site^21^.

Thus, these data raise the possibility that NB6 may be broadly effective in rescuing other class II mutants.

The opposite defect, a right shift induced by class III mutations, was corrected by using the cAMP antagonist TRIP8b_nano_ in variants (wt/M153I and wt/E246K) that are unrelated to cAMP binding defects. In these cases, TRIP8b_nano_ can compensate for the defect rather than directly rescue the underlying molecular alteration. Whether TRIP8b_nano_ can exert its effect in neurons remains an open question. Neurons expressing HCN1, also express the full-length protein TRIP8b, from which TRIP8b_nano_ was derived, raising the possibility of competition between the two^22^.

In search of alternatives to reduce the current at physiologically relevant voltages in class III and class IV variants, we tested other known HCN1 inhibitors. GOF mutations that increase instantaneous current could be counteracted by ivabradine, the only HCN-specific drug currently approved for clinical use. However, despite this theoretical suitability, ivabradine shows little blood-brain barrier permeability,^24^ limiting its use for the treatment of neuronal HCN1 dysfunction. An alternative treatment may be provided by newly available brain permeant compounds such as J&J12e. In our proof-of-concept experiments we showed that J&J12e is indeed able to rescue class III mutants at low nanomolar concentrations (25-50 nM) within a physiologically relevant range of membrane potentials (-90 to -40 mV).

### Functional variant analysis as a predictor of clinical phenotype and disease severity

In this international cohort of individuals carrying *HCN1* variants, we identified a striking relationship between HCN1 functional effect and clinical expression. LOF variants were markedly enriched among probands without epilepsy, whereas non-LOF variants including GOF, mixed, and wild-type-like variants were predominantly associated with epilepsy. This association remained significant in logistic regression analysis, indicating substantially reduced odds of epilepsy among LOF carriers within this cohort.

Beyond epilepsy occurrence, the type of functional alteration also segregated with clinical severity. Among individuals with epilepsy, LOF variants clustered toward milder phenotypes, whereas severe neonatal and infantile onset DEE were confined to the non-LOF group. The severity gradient was highly significant and driven largely by GOF variants, suggesting that increased or dysregulated HCN1 channel activity may confer greater vulnerability during early neurodevelopment.

These findings support a model in which the type of channel dysfunction, not merely its presence, critically shapes disease presentation. *HCN1* channels regulate intrinsic neuronal excitability and dendritic integration, particularly in cortical and hippocampal pyramidal neurons. Gain-of-function or persistent channel activation may enhance aberrant depolarizing currents during critical developmental windows, promoting network hyperexcitability and impairing circuit maturation, thereby predisposing to DEE. In contrast, reduced *HCN1* function may alter oscillatory dynamics and thalamocortical synchrony without necessarily driving catastrophic early epileptogenesis, instead favouring generalized or absence-spectrum phenotypes.

When epilepsy syndromes were grouped according to ILAE criteria, functional class was again strongly associated with phenotype distribution. DEE occurred exclusively in the non-LOF group, whereas generalized/absence-spectrum epilepsies were enriched among LOF carriers. Although these analyses are limited by modest sample size and sparsity of some categories, the magnitude and consistency of the associations suggest that functional direction is a major determinant of clinical trajectory in *HCN1*-related disease.

Taken together, these findings indicate that *HCN1*-related disorders stratify along a biologically meaningful axis defined by the direction of channel dysfunction. This functional stratification may have implications for prognostic counselling and for future therapeutic strategies aimed at restoring physiological channel activity.

## Conclusions

Beyond advancing the understanding of HCN1 channel biophysics, our findings provide a clinically meaningful framework for variant interpretation. In several proof-of-concept experiments we demonstrate that available allosteric modulators can restore wt-like properties in mutants from three distinct functional classes.

Our systematic comparison of mutant classes and topological localization further revealed an unexpected subgroup of LOF variants identified in patients without epilepsy. These results highlight the complexity of HCN1-related dysfunction and emphasize the value of an integrated classification system for both mechanistic insight and therapeutic development. By showing that the direction of channel dysfunction correlates with epilepsy risk and severity, we propose a model that may assist prognostic assessment and guide future precision therapeutic strategies in HCN1-related disease.

## Data availability

The authors declare that the data supporting the findings of this study are available within the article and its Supplementary Information file, and from the corresponding authors upon request. Previously published PDB codes are: 6uqf.

## Acknowledgements

We thank Dr. Benjamin Hall (Lundbeck A/S, Denmark) for providing J&J12e and Org-34167.

## Supplementary material

Supplementary material is available at *Brain* online. This includes Supplementary Figures, Tables and Methods files.

## References

1. Santoro B, Shah MM. Hyperpolarization-Activated Cyclic Nucleotide-Gated Channels as Drug Targets for Neurological Disorders. Annu Rev Pharmacol Toxicol. 2020;60(1):109–131. doi:10.1146/annurev-pharmtox-010919-023356

2. DiFrancesco D. Funny channel gene mutations associated with arrhythmias. J Physiol. 2013;591(17):4117–4124. doi:10.1113/jphysiol.2013.253765

3. Emery EC, Young GT, McNaughton PA. HCN2 ion channels: an emerging role as the pacemakers of pain. Trends Pharmacol Sci. 2012;33(8):456–463. doi:10.1016/j.tips.2012.04.004

4. Marini C, Porro A, Rastetter A, et al. *HCN1* mutation spectrum: from neonatal epileptic encephalopathy to benign generalized epilepsy and beyond. Brain. 2018;141(11):3160–3178. doi:10.1093/brain/awy263

5. Houdayer C, Phillips AM, Chabbert M, et al. *HCN2* -Associated Neurodevelopmental Disorders: Data from Patients and *Xenopus* Cell Models. Ann Neurol. 2025;98(3):573–589. doi:10.1002/ana.27277

6. Nava C, Dalle C, Rastetter A, et al. De novo mutations in HCN1 cause early infantile epileptic encephalopathy. Nat Genet. 2014;46(6):640–645. doi:10.1038/ng.2952

7. McKenzie CE, Forster IC, Soh MS, et al. Cation leak: a common functional defect causing *HCN1* developmental and epileptic encephalopathy. Brain Commun. 2023;5(3). doi:10.1093/braincomms/fcad156

8. Xie C, Liu F, He H, et al. Novel HCN1 Mutations Associated With Epilepsy and Impacts on Neuronal Excitability. Front Mol Neurosci. 2022;15. doi:10.3389/fnmol.2022.870182

9. Porro A, Abbandonato G, Veronesi V, et al. Do the functional properties of HCN1 mutants correlate with the clinical features in epileptic patients? Prog Biophys Mol Biol. 2021;166:147–155. doi:10.1016/j.pbiomolbio.2021.07.008

10. Bleakley LE, McKenzie CE, Soh MS, et al. Cation leak underlies neuronal excitability in an HCN1 developmental and epileptic encephalopathy. Brain. 2021;144(7):2060–2073. doi:10.1093/brain/awab145

11. Lee CH, MacKinnon R. Voltage Sensor Movements during Hyperpolarization in the HCN Channel. Cell. 2019;179(7):1582–1589.e7. doi:10.1016/j.cell.2019.11.006

12. Burtscher V, Mount J, Huang J, et al. Structural basis for hyperpolarization-dependent opening of human HCN1 channel. Nat Commun. 2024;15(1):5216. doi:10.1038/s41467-024-49599-x

13. Lee CH, MacKinnon R. Structures of the Human HCN1 Hyperpolarization-Activated Channel. Cell. 2017;168(1-2):111–120.e11. doi:10.1016/j.cell.2016.12.023

14. Tao X, Lee A, Limapichat W, Dougherty DA, MacKinnon R. A Gating Charge Transfer Center in Voltage Sensors. Science (1979). 2010;328(5974):67–73. doi:10.1126/science.1185954

15. Decher N, Chen J, Sanguinetti MC. Voltage-dependent Gating of Hyperpolarization-activated, Cyclic Nucleotide-gated Pacemaker Channels. Journal of Biological Chemistry. 2004;279(14):13859–13865. doi:10.1074/jbc.M313704200

16. Schmidpeter PAM, Wu D, Rheinberger J, et al. Anionic lipids unlock the gates of select ion channels in the pacemaker family. Nat Struct Mol Biol. 2022;29(11):1092–1100. doi:10.1038/s41594-022-00851-2

17. Zhou L, Siegelbaum SA. Gating of HCN Channels by Cyclic Nucleotides: Residue Contacts that Underlie Ligand Binding, Selectivity, and Efficacy. Structure. 2007;15(6):655–670. doi:10.1016/j.str.2007.04.012

18. Pfleger C, Kusch J, Kondapuram M, et al. Allosteric signaling in C-linker and cyclic nucleotide-binding domain of HCN2 channels. Biophys J. 2021;120(5):950–963. doi:10.1016/j.bpj.2021.01.017

19. Saponaro A, Bauer D, Giese MH, et al. Gating movements and ion permeation in HCN4 pacemaker channels. Mol Cell. 2021;81(14):2929–2943.e6. doi:10.1016/j.molcel.2021.05.033

20. Bassetto CAZ, Costa F, Guardiani C, Bezanilla F, Giacomello A. Noncanonical electromechanical coupling paths in cardiac hERG potassium channel. Nat Commun. 2023;14(1):1110. doi:10.1038/s41467-023-36730-7

21. Sharifzadeh AS, Castelli R, Porro A, et al. Extracellular activation of HCN4 by a subtype-specific nanobody. Nat Commun. 2025;16(1):10804. doi:10.1038/s41467-025-65852-3

22. Saponaro A, Cantini F, Porro A, et al. A synthetic peptide that prevents cAMP regulation in mammalian hyperpolarization-activated cyclic nucleotide-gated (HCN) channels. Elife. 2018;7. doi:10.7554/eLife.35753

23. 23. European Medicines Agency (EMA) Product Information Ivabradine [Internet] Amsterdam: European Medicines Agency; 2018. [Cited 2019 Aug 10]. Available from Https://Www.Ema.Europa.Eu/En/Documents/Product-Information/Procoralan-Epar-Product-Information_en.Pdf.

24. Savelieva I, Camm AJ. Novel If Current Inhibitor Ivabradine: Safety Considerations. In: Heart Rate Slowing by I_f_ Current Inhibition. KARGER; 2006:79–96. doi:10.1159/000095430

25. Saponaro A, Krumbach JH, Chaves-Sanjuan A, et al. Structural determinants of ivabradine block of the open pore of HCN4. Proceedings of the National Academy of Sciences. 2024;121(27). doi:10.1073/pnas.2402259121

26. McKenzie CE, Hung A, Phillips AM, Soh MS, Reid CA, Forster IC. The Potential Antidepressant Compound Org 34167 Modulates HCN Channels Via a Novel Mode of Action. Mol Pharmacol. 2023;104(2):62–72. doi:10.1124/molpharm.123.000676

27. Bleakley LE, McKenzie CE, Aung KP, et al. Org 34167 rescues voltage dependence of mutant channels and normalizes hyperexcitability in *HCN1* epilepsy. Epilepsia. 2025;66(12):5082–5095. doi:10.1111/epi.18585

28. Hill WG, Southern NM, MacIver B, et al. Isolation and characterization of the *Xenopus* oocyte plasma membrane: a new method for studying activity of water and solute transporters. American Journal of Physiology-Renal Physiology. 2005;289(1):F217–F224. doi:10.1152/ajprenal.00022.2005

29. Handlin LJ, Dai G. Direct regulation of the voltage sensor of HCN channels by membrane lipid compartmentalization. Nat Commun. 2023;14(1):6595. doi:10.1038/s41467-023-42363-7

30. McClure KJ, Maher M, Wu N, et al. Discovery of a novel series of selective HCN1 blockers. Bioorg Med Chem Lett. 2011;21(18):5197–5201. doi:10.1016/j.bmcl.2011.07.051

31. Harde E, Hierl M, Weber M, et al. Selective and brain-penetrant HCN1 inhibitors reveal links between synaptic integration, cortical function, and working memory. Cell Chem Biol. 2024;31(3):577–592.e23. doi:10.1016/j.chembiol.2023.11.004

32. Merseburg A, Kasemir J, Buss EW, et al. Seizures, behavioral deficits and adverse drug responses in two new genetic mouse models of HCN1 epileptic encephalopathy. Elife. 2022;11. doi:10.7554/eLife.70826

33. Bleakley LE, McKenzie CE, Soh MS, et al. Cation leak underlies neuronal excitability in an HCN1 developmental and epileptic encephalopathy. Brain. 2021;144(7):2060–2073. doi:10.1093/brain/awab145

34. Upadhyay K, Nigam N, Gupta S, Tripathi SK, Jain A, Puri B. Current and future therapeutic approaches of CFTR and airway dysbiosis in an era of personalized medicine. J Family Med Prim Care. 2024;13(6):2200–2208. doi:10.4103/jfmpc.jfmpc_1085_23

35. Marson FAL, Bertuzzo CS, Ribeiro JD. Classification of CFTR mutation classes. Lancet Respir Med. 2016;4(8):e37–e38. doi:10.1016/S2213-2600(16)30188-6

36. DiFrancesco D, Tortora P. Direct activation of cardiac pacemaker channels by intracellular cyclic AMP. Nature. 1991;351(6322):145–147. doi:10.1038/351145a0

37. Porro A, Saponaro A, Gasparri F, et al. The HCN domain couples voltage gating and cAMP response in hyperpolarization-activated cyclic nucleotide-gated channels. Elife. 2019;8. doi:10.7554/eLife.49672

38. Porro A, Saponaro A, Castelli R, et al. A high affinity switch for cAMP in the HCN pacemaker channels. Nat Commun. 2024;15(1):843. doi:10.1038/s41467-024-45136-y

39. Tao X, Lee A, Limapichat W, Dougherty DA, MacKinnon R. A Gating Charge Transfer Center in Voltage Sensors. Science (1979). 2010;328(5974):67–73. doi:10.1126/science.1185954

40. Lacroix JJ, Labro AJ, Bezanilla F. Properties of Deactivation Gating Currents in Shaker Channels. Biophys J. 2011;100(5):L28–L30. doi:10.1016/j.bpj.2011.01.043

