## Supplementary Material for "Comprehensive classification of HCN1 variants linked to neurodevelopmental disorders with and without epilepsy"

**Castelli et al.,**

### Supplementary Methods

#### Constructs

The cDNA encoding full-length human HCN1 (hHCN1), naked or fused in frame with either eGFP or TagRFP at the N-terminus, hERG and $Na_v1.5$ were previously cloned into the pcDNA 3.1 (Invitrogen) mammalian expression vector. Single point mutations were introduced using the QuickChange XL-II kit (Agilent Technologies) and all constructs were sequenced to ensure that no additional variants had been introduced. Reported mutations located in the distal C-terminus of HCN1 were not characterized in this study, as a delta C-term construct (P638*) displayed no detectable functional defects when measured in our experimental conditions.

#### HEK293T and HEK293F cells culture and transfection

HEK293T cells were cultured in Dulbecco's modified Eagle's medium (Euroclone), while HEK293F were grown in Freestyle medium (Thermo Fisher), both supplemented with 10% fetal bovine serum (Euroclone), 1% Pen Strep (100 U/ml of penicillin and 100 μg/ml of streptomycin) and grown at 37 °C with 5% $CO_2$. For electrophysiology experiments, HEK293T cells were transiently transfected with 1 μg of the HCN1-containing plasmid (homotetramers) or 0.5 μg of each plasmid (wild-type and mutant, heterotetramers) along with 0.3 μg of EGFP-containing vector (pmaxGFP, Amaxa Biosystems) to track for positive cell transfection, using Turbofect transfection reagent (Thermo Fisher Scientific, Germany). Cells were periodically tested for Mycoplasma contamination using MycoAlter detection kit (Lonza) and always resulted negative.

#### Patch clamp recordings in HEK cells

24 h after transfection HEK293T or HEK293F cells were dispersed and single $GFP^+$ cells were selected for patch-clamp recordings. Each set of experiments contains data from controls and mutants measured on the same day. Currents were recorded in whole-cell configuration at room temperature, either with an ePatch amplifier (Elements, Cesena, Italy) or with an Axopatch 200b amplifier (Molecular Devices); data acquired with the Axopatch 200b amplifier were digitized with an Axon Digidata 1550B (Molecular Devices) converter. Signals were acquired with a sampling rate of 5 kHz and low pass filtered at 2.5 kHz. Data analysis was performed using Clampfit 10.7 (Molecular devices). Patch pipettes were pulled from 1.5 mm O.D. and

0.86 mm I.D. borosilicate glass capillaries (Sutter, Novato, CA) and had resistances ranging from 3 to 6 MΩ. For HCN1 channels recordings, patch pipettes were filled with a solution containing 10 mM NaCl, 130 mM KCl, 1 mM egtazic acid (EGTA), 0.5 mM MgCl2, 2 mM ATP (magnesium salt), and 5 mM HEPES–KOH buffer (pH 7.2), while the extracellular bath solution contained 110 mM NaCl, 30 mM KCl, 1.8 mM CaCl2, 0.5 mM MgCl2, and 5 mM HEPES–KOH buffer (pH 7.4). To record HEK293F endogenous currents patch pipettes were filled with a solution containing 155 mM KCl, 3 mM MgCl2, 1 mM EGTA, 1 mM ATP (magnesium salt), 10 mM HEPES-KOH buffer (pH 7.2), while the extracellular solution contained 150 mM NaCl, 5 mM KCl, 1 mM CaCl2, 3 mM MgCl2 and 10 mM HEPES-NaOH buffer (pH 7.4). For hERG channel recordings patch pipettes were filled with a solution containing 130 mM KCl, 1 mM MgCl2, 5 mM EGTA, 5 mM ATP (magnesium salt) and 10 mM HEPES-KOH buffer (pH 7.2), while the extracellular bath solution contained 140 mM NaCl, 4 mM KCl, 2.5 mM CaCl2, 10 mM Glucose, 5 mM HEPES-NaOH buffer (pH 7.45). For $Na_v1.5$ channel recordings patch pipettes were filled with a solution containing 5 mM NaCl, 140 mM CsCl, 2 mM MgCl2, 5 mM EGTA, 4 mM ATP (magnesium salt) and 10 mM HEPES-CsOH buffer (pH 7.4), while the extracellular bath solution contained 135 mM NaCl, 4 mM KCl, 1 mM CaCl2, 2 mM MgCl2, 20 mM Glucose, 10 mM HEPES-NaOH buffer (pH 7.4).

Where indicated, NB6 was added to the extracellular bath solution to reach a final concentration of 20 μM. Controls were treated with the nanobody buffer which contains: 150 mM NaCl, 20 mM HEPES and 10% (w/v) Glycerol at pH 7.5 supplemented with 1:1000 cOmplete$^{TM}$, EDTA free (Sigma-Aldrich). Both buffer and nanobodies were stored at -80 °C until the day of the experiment where single-use aliquots were slowly thawed in ice. Ivabradine block was studied by superfusing the drug at 30 μM during repetitive application of activating (-110 mV; 1.5 s) and deactivating (+5 mV, 0.5 s) voltage steps every 5.5 seconds, from a holding potential of -20 mV, until development of the full block. Fractional current block was calculated as the ratio between the steady state current at the full block and the steady state current measured at -110 mV, just before drug application. $TRIP8b_{nano}$ was produced as a synthetic peptide (Caslo, Denmark) and later dissolved in milliQ water to make 10 mM stock solutions. Single-use aliquots were stored at -20°C until the day of the experiment, where they were diluted to 10 μM in the intracellular recording solution. J&J12e and Org-34617 compounds (provided by H. Lundbeck A/S, Denmark) were dissolved in DMSO to make 10 mM stock solutions. Single-use aliquots were prepared and stored at -20°C until the day of the experiment. From the stock solution, all compounds were diluted to the desired concentration and kept in the extracellular recording solution throughout the entire patch clamp session; controls were treated with the vehicle (DMSO).

To assess HCN1 channel activation curves, different voltage-clamp protocols were applied. For HCN1 holding potential was -20 mV (1 s), with steps from -30 mV to -120 mV (-10 mV 26 increments, 3.5 s) and tail currents recorded at -40 mV (3.5 s). For hERG holding potential was -80 mV (1s), with steps from -60 mV to +60 mV (Δ of +10 mV, 4 s) and tail currents were collected at -50 mV (6 s). For non-transfected HEK293F cells (NT), holding was -80 mV (100 ms), with steps from -100 mV to +100 mV (300 ms), and return to the holding potential of -80 mV (100 ms). For $Na_v1.5$ holding potential was -120 mV (100 ms), followed by steps from 100 mV to +85 mV (Δ of +5 mV, 100 ms), and return to the holding potential of -100 mV (200 ms).

Only cells in which a 1 GΩ seal or better was achieved were kept for analysis. For I/V plots currents were normalized to cell capacitance, indicated as $I_{SS}$ (pA/pF). Neither series resistance compensation nor leak correction were applied.

Mean activation curves were obtained by fitting maximal tail current amplitude, plotted against the preconditioning voltage step, with the Boltzmann equation: $y = 1/[1+\exp((V-V_{1/2})/k)]$, where V is voltage, y the fractional activation, $V_{1/2}$ the half-activation voltage, and k the inverse slope factor in mV ($k = -RT/zF$). Mean activation curves were obtained by fitting individual curves from each cell to the Boltzmann equation and then averaging all the obtained values.

Data were analyzed with Clampfit (Molecular devices) and Origin (OriginLab) softwares and are presented as mean ± SEM. Statistical analysis was performed with the Student's t-test for unpaired data or One-way ANOVA with Fisher's test, as indicated for each experiment. Significance level was set to $p = 0.05$.

**Confocal fluorescence imaging**

To study homotetrameric channel assemblies, HEK293F cells were transfected with eGFP-hHCN1 fusion constructs (wild-type or mutant) and, 24 h after transfection, incubated for 5 minutes with CellMask™ red, a plasma membrane dye. To study heterotetrameric assemblies, cells were co-transfected with wild-type TagRFP-hHCN1 (red) and mutant eGFP-hHCN1(green) in a 1:1 ratio. Confocal imaging of HEK293F cells was carried out 24 h after transfection using a Nikon Eclipse-Ti inverted confocal microscope interface with an A1 series of confocal laser point scanning system which allows for excitation at 405, 488, 561 and 640 nm. Samples plated on 35 mm glass Petri dishes were observed with a 60x 1.4 NA oil immersion objective (Nikon System). The pinhole aperture was set to 1.0 Airy. Images were collected using low excitation power (488, 561 and 640 nm) and acquiring the emission range through bandpass filters 525/50, 595/50 and 700/70 for eGFP, TagRFP and CellMask™ Plasma Membrane Stain, respectively, by means of the built-in GaAsP PMT detectors of the confocal microscope. We generated intensity profiles of the signal at the plasma membrane by measuring the fluorescence intensities of the red (CellMaskTM or wild-type TagRFP-hHCN1) and green (wild-type or mutant eGFP-hHCN1) signals encompassing the membrane. These intensities were normalized [0;1] and plotted as a function of the pixels which indicate the measured distance across the membrane. Overlapping peaks indicate that the two signals colocalize. Channel membrane localization was measured as the ratio ($F_{PR}/F_{PM}$) between the fluorescence signal at the periplasmic ring ($F_{PR}$) over the signal at the plasma membrane ($F_{PM}$). For each cell, the $F_{PR}/F_{PM}$ value was determined as the average ratio between 10 points at the periplasmic ring and plasma membrane, each. Imaging data were analyzed using ImageJ (U.S. National Institutes of Health, Bethesda, MD, USA) and Origin (OriginLab) softwares. All data are presented as mean ± S.E.M. Statistical comparisons were made with One-Way ANOVA test with a significance threshold of $p = 0.05$.

**Analysis of clinical phenotypes**

**Patients and variant selection**

Probands carrying *HCN1* variants, either novel or previously published were included. Variants were ascertained through an international collaboration involving centres in Italy, United Kingdom, France, Netherlands, Poland, Switzerland, Ireland, Germany, United States of America, and Australia. Both de novo and familial cases were eligible. Variants were collected from participating clinicians and/or existing literature and were annotated using the reference transcript NM_021072.4. Variant nomenclature follows HGVS recommendations. When available, segregation data and inheritance were recorded. Overall, the study cohort consisted of 49 patients carrying HCN variants who met the inclusion criteria and for whom clinical data were available.

#### Clinical data collection and phenotypic classification

For each proband, standardized clinical data were collected from medical records and clinician reports. Variables of interest included:

- epilepsy status (present/absent);
- age at seizure onset;
- epilepsy syndrome and seizure types classified according to ILAE criteria when possible;
- epilepsy severity category;
- neurodevelopmental outcome, including developmental delay/intellectual disability, language impairment, autistic traits, and behavioural disturbances;
- additional neurological features and/or dysmorphisms;
- brain MRI findings (normal/abnormal and main reported abnormalities).

Epilepsy severity was categorized a priori as severe, moderate, or mild. Severe epilepsy was defined as neonatal- or infantile-onset developmental and epileptic encephalopathy (DEE), including drug-resistant DEE. Mild epilepsy included generalized genetic epilepsies (GGE), febrile seizures plus (FS+), GEFS+, and focal epilepsies without encephalopathy. Moderate severity included unclassified epilepsies, focal hemiclonic seizures, or a single episode of refractory status epilepticus without a sustained DEE course. In select cases, specifically patient ID#14 (S272P), patient ID#40 (G391C) and patient ID#48 (R548H), the severity classification was attuned based on clinical judgement integrating seizure frequency and overall developmental profile as reported in the medical records.

#### Statistical Analysis (clinical data)

Associations between functional class and clinical phenotypes were evaluated as follows. For genotype-phenotype correlation analyses, variants were grouped into two categories: loss-of-function (LOF; Classes I–II) and non-LOF (Classes III-IV, including gain-of-function [GOF], mixed LOF/GOF, and wild-type–like variants).

*Epilepsy risk:* Epilepsy occurrence (present/absent) was treated as a binary outcome. Associations between functional class and epilepsy status were first assessed using Fisher's exact test due to small cell counts. Univariable logistic regression was then performed to estimate odds ratios (ORs) and 95% confidence intervals (CIs), with epilepsy status as the dependent variable and functional class (LOF versus non-LOF) as the predictor. Given the limited sample size and imbalance between groups, ORs were interpreted cautiously.

*Epilepsy severity:* Severity analyses were restricted to probands with epilepsy. Epilepsy severity was categorized a priori as mild, intermediate, or severe, as defined above. The association between functional class and severity distribution was assessed using the Fisher–Freeman–Halton exact test for r×c contingency tables. To further quantify the effect on severe phenotypes, severity was dichotomized (severe versus non-severe) and analysed using Fisher's exact test and logistic regression to estimate ORs and 95% CIs. Because severe phenotypes were absent in the LOF group, effect estimates were interpreted in the context of complete separation.

*ILAE epilepsy/syndrome categories:* Among probands with epilepsy, the type of epilepsies and syndromes were grouped according to ILAE classification (DEE, GGE/absence-spectrum, focal epilepsies, FS+/GEFS+, other/unclassified). Associations between functional class and syndrome distribution were assessed using the Fisher–Freeman–Halton exact test.

All tests were two-sided, and p-values $<0.05$ were considered statistically significant.

### Supplementary Figures

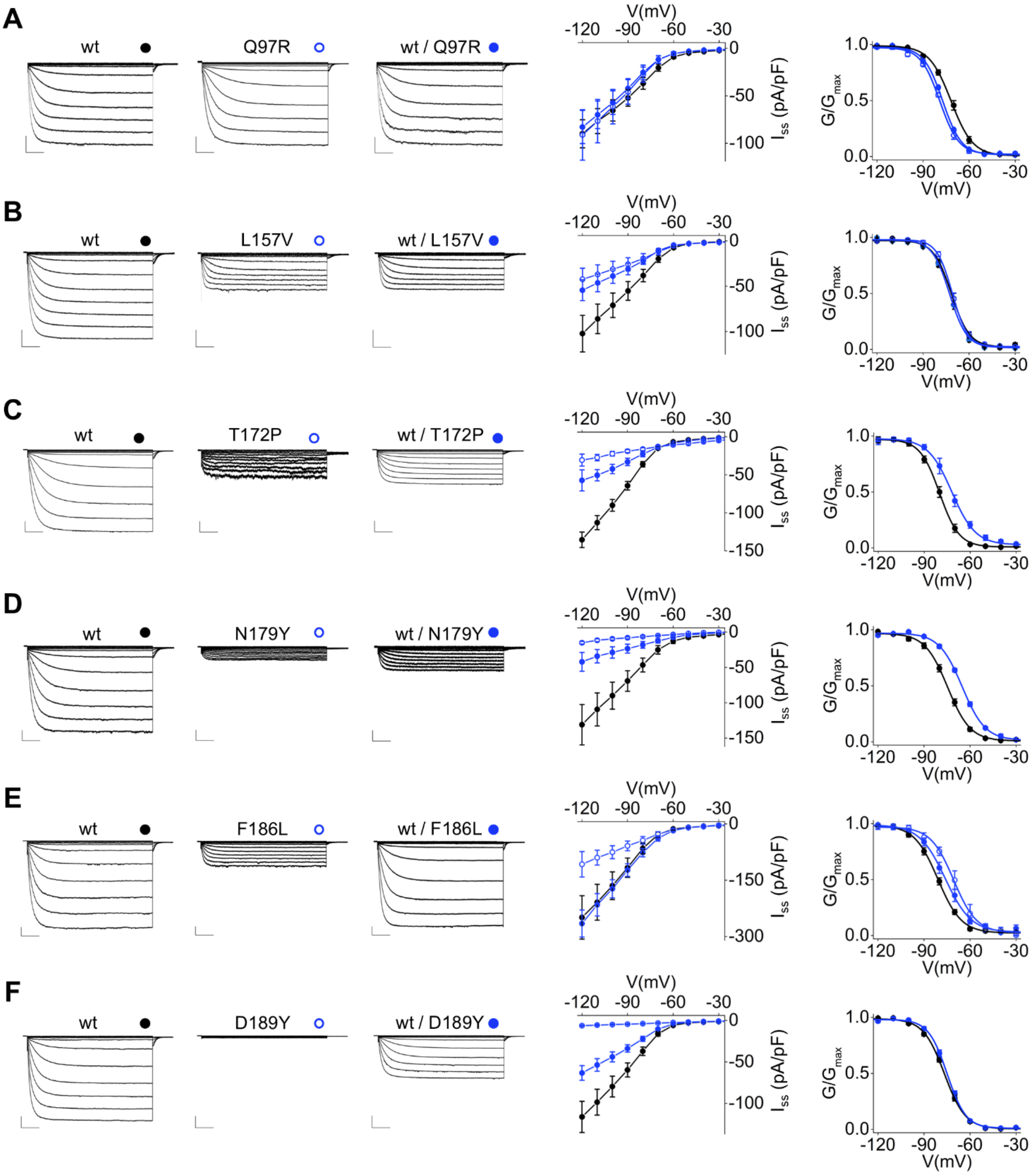

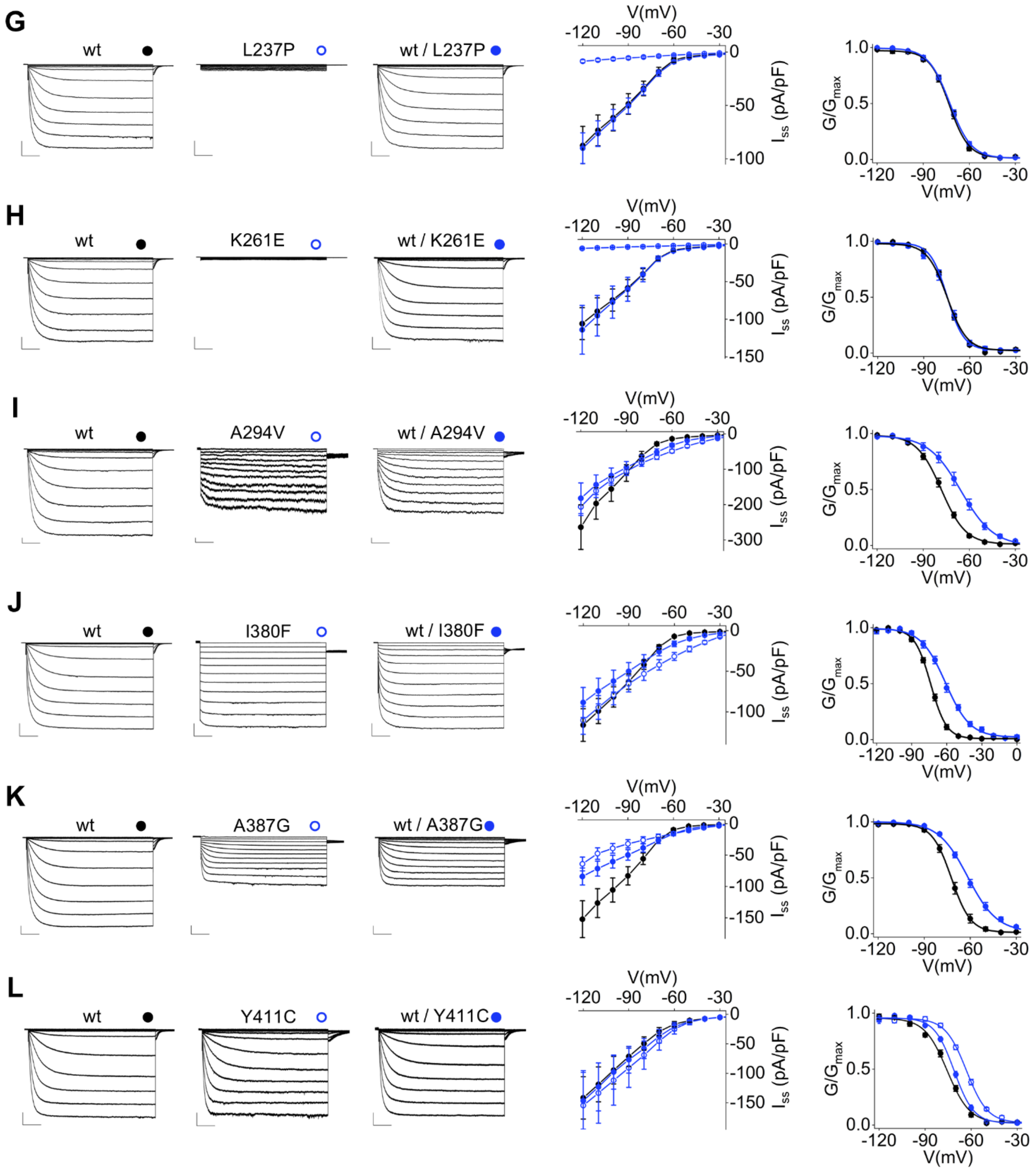
G
wt
L237P
wt / L237P
H
wt
K261E
wt / K261E
I
wt
A294V
wt / A294V
J
wt
I380F
wt / I380F
K
wt
A387G
wt / A387G
L
wt
Y411C
wt / Y411C
V(mV)
$I_{ss}$ (pA/pF)
$G/G_{max}$

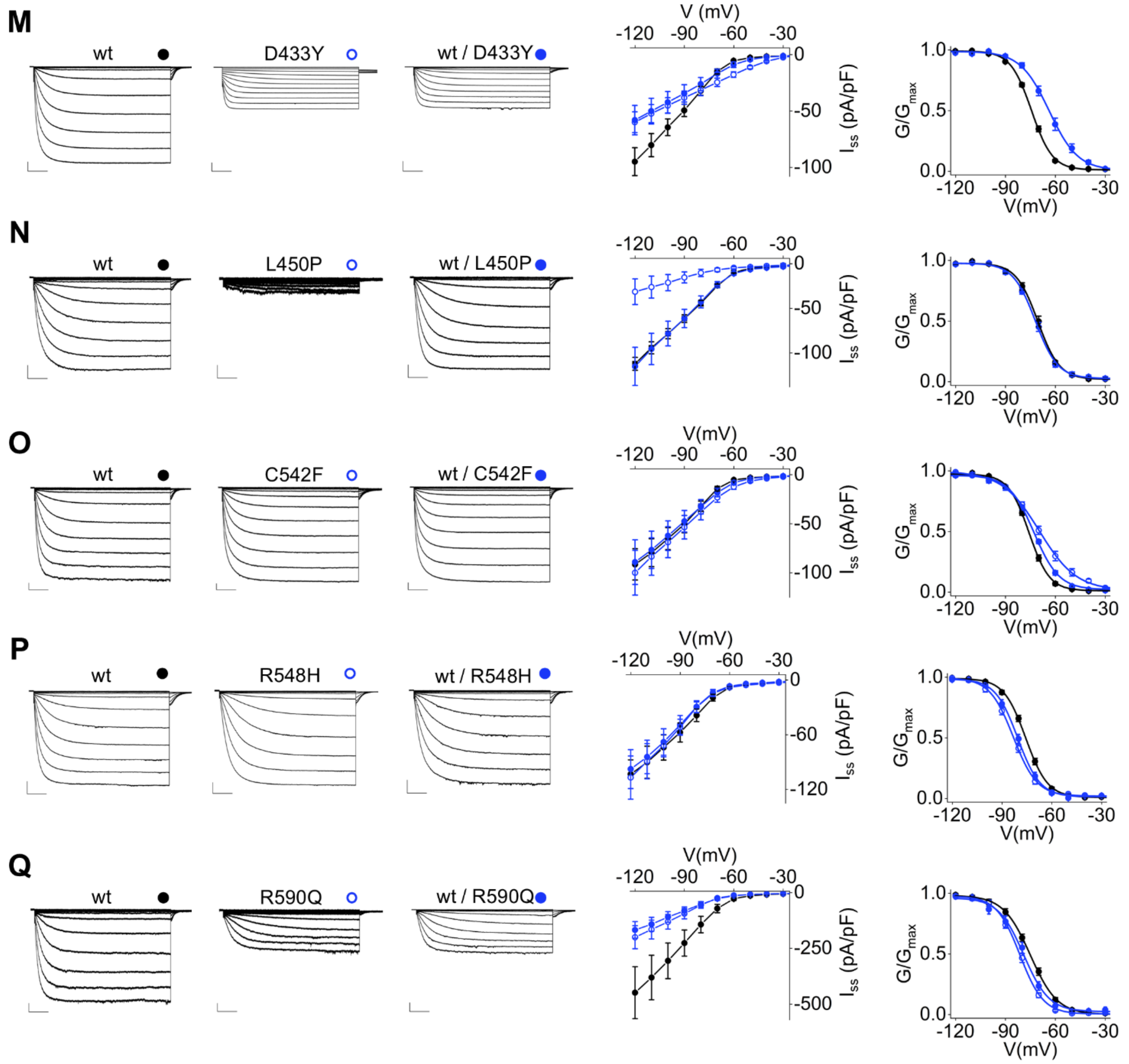


**Supplementary Figure 1a, 1b and 1c. Functional properties of HCN1 variants.** Representative whole-cell currents of HCN1 mutants recorded in HEK293 cells: Q97R (**A**), L157V (**B**), T172P (**C**), N179Y (**D**), F186L (**E**), D189Y (**F**), L237P (**G**), K261E (**H**), A294V (**I**), I380F (**J**), A387G (**K**), Y411C (**L**), D433Y (**M**), L450P (**N**), C542F (**O**), R548H (**P**) and R590Q (**Q**). Cells were transiently transfected with HCN1 wild type alone (wt, black dot), HCN1 mutant alone (homotetramers, empty blue dot) or both (heterotetramers, solid blue dot). Traces shown from -20 mV to -120 mV. Scale bars: 250 pA and 500 ms. Middle: corresponding mean I/V relationships with current density at steady-state ($I_{ss}$, pA/pF). Data are mean ± SEM. Right: mean activation curves obtained from HCN1 wt alone (black dots), HCN1 mutant alone (empty blue dots) or both (solid blue dots). Data fit to the Boltzmann equation are plotted as solid lines. Data points are mean ± SEM. $I_{ss}$ values measured at -120 mV, $V_{1/2}$, $\Delta V_{1/2}$, inverse slope factors (k) and number of cells (n) for each experiment shown are reported in Supplementary Tables 1 and 2 along with the details on statistical analysis.

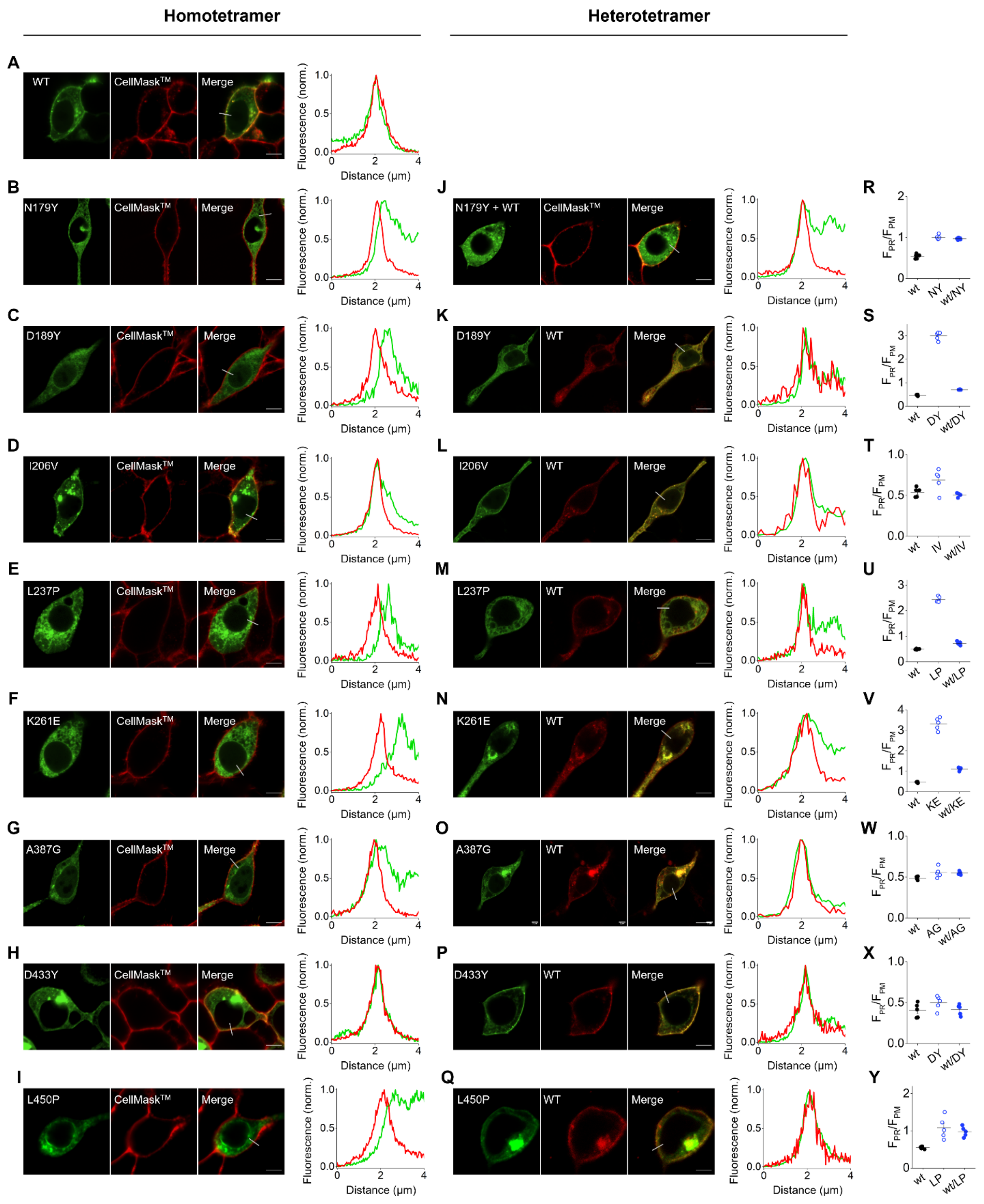


**Supplementary Figure 2. Confocal microscopy analysis of HCN1 variants.** Confocal images of mutant eGFP-HCN1 expressed alone or co-expressed with TagRFP-HCN1 wt in HEK293F cells. Homotetramers, from left to right: eGFP-HCN1 wt (**A**), N179Y (**B**), D189Y (**C**), I206V (**D**), L237P (**E**), K261E (**F**), A387G (**G**), D433Y (**H**) or L450P (**I**) (green), CellMask$^{TM}$ plasma membrane dye (red) and merge of the two signals (yellow). Scale bar 5 µm. Right: representative normalized intensity profiles of fluorescent signals encompassing the plasma membrane (grey line drawn on the merged image). Heterotetramers, from left to right: eGFP-HCN1 N179Y (**J**), D189Y (**K**), I206V (**L**), L237P (**M**), K261E (**N**), A387G (**O**), D433Y (**P**) or L450P (**Q**) (green), TagRFP-HCN1 wt (red) and merge of the two signals (yellow). Right: normalized fluorescence intensity profiles obtained by plotting all points encompassed by the grey line drawn on the merged image. Scale bar 5 μm. Overlapping peaks indicate plasma membrane co-localization. (**R-Y**) Mean fluorescence signal intensity expressed as $F_{PR}/F_{PM}$ ratio. Data shown as mean ± SEM. Quantitative analysis of $F_{PR}/F_{PM}$ ratio values, mean Pearson correlation coefficients (r), peak distance (PD) values and number of cells (n) are reported in Supplementary Tables 3 and 4 along with the details on statistical analysis.

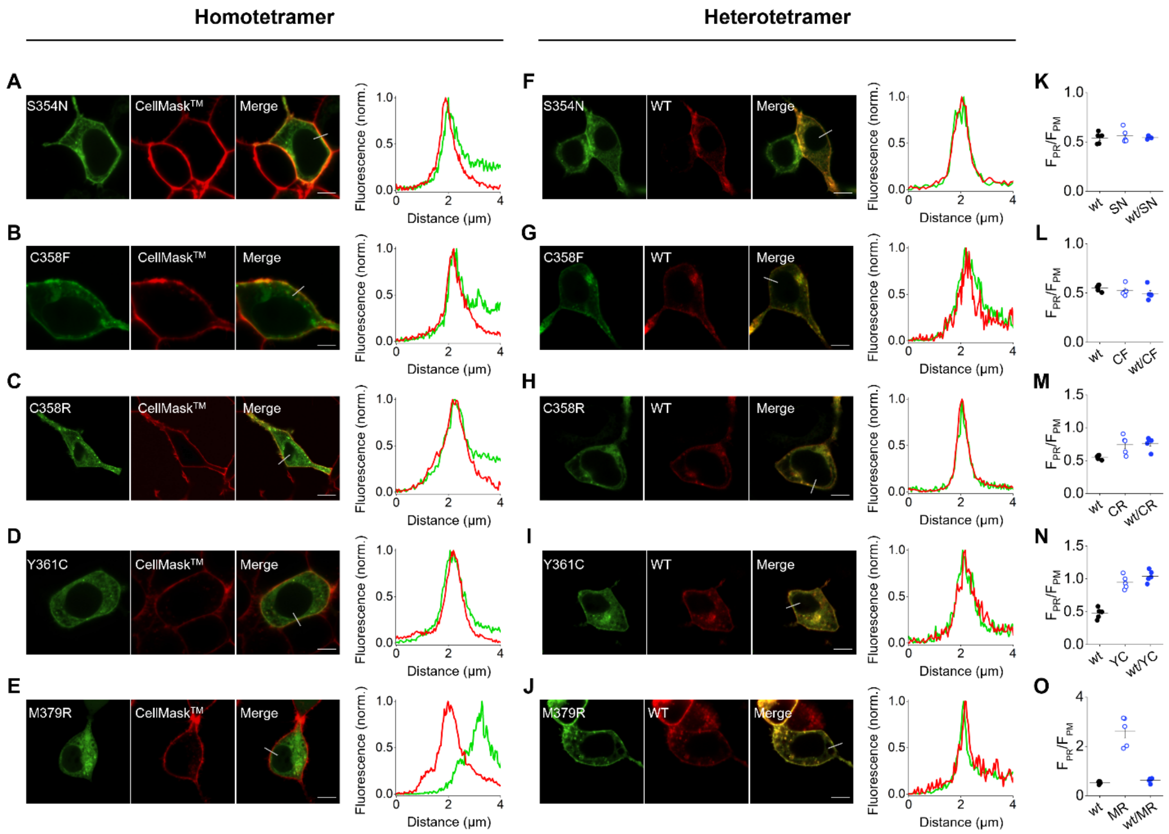


**Supplementary Figure 3. Confocal microscopy analysis of LOF HCN1 variants linked to ND.** Confocal images of non-epileptic mutant eGFP-HCN1 expressed alone or co-expressed with TagRFP HCN1 wt in HEK293F cells. Homotetramers, from left to right: eGFP-HCN1 S354N (**A**), C358F (**B**), C358R (**C**), Y361C (**D**) or M379R (**E**) (green), CellMask$^{TM}$ plasma membrane dye (red) and merge of the two signals (yellow). Scale bar 5 µm. Right: representative normalized intensity profiles of fluorescent signals encompassing the plasma membrane (grey line drawn on the merged image). Heterotetramers, from left to right: eGFP HCN1 S354N (**F**), C358F (**G**), C358R (**H**), Y361C (**I**) or M379R (**J**) (green), TagRFP-HCN1 wt (red) and merge of the two signals (yellow). Right: normalized fluorescence intensity profiles obtained by plotting all points encompassed by the grey line drawn on the merged image. Scale bar 5 µm. Overlapping peaks indicate plasma membrane co-localization. (**K-O**) Mean fluorescence signal intensity expressed as $F_{PR}/F_{PM}$ ratio. Data shown as mean ± SEM. Quantitative analysis of $F_{PR}/F_{PM}$ ratio values, mean Pearson correlation coefficients (r), peak distance (PD) values and number of cells (n) are reported in Supplementary Tables 3 and 4 along with the details on statistical analysis.

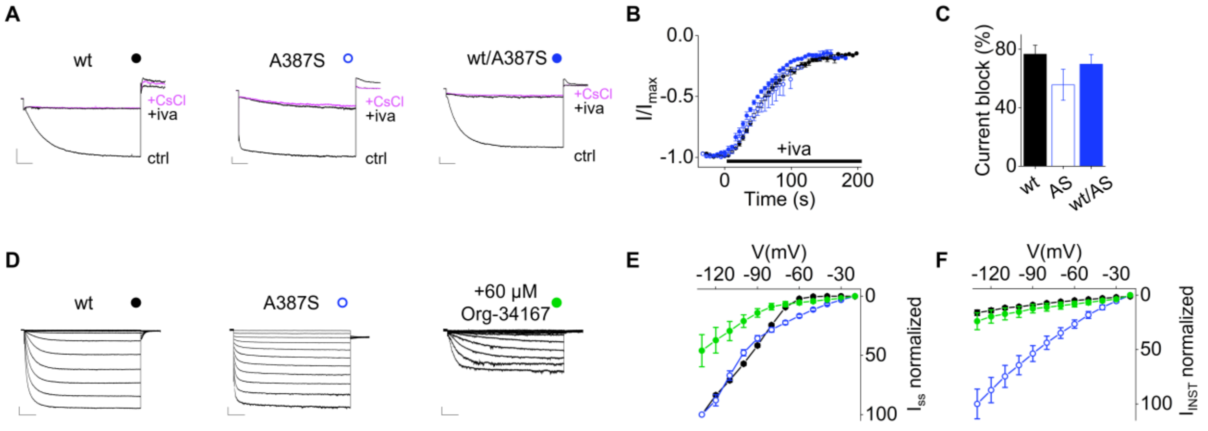


**Supplementary Figure 4. Effect of ivabradine and Org-34167 on HCN1 wt and A387S.** (**A**) Representative current traces measured at -110 mV of HCN1 wt (left), A387S (center) and wt/A387S (right), in control solution (ctrl) and after superfusion of 30 µM ivabradine (+iva) at steady-state block, followed by addition of 5 mM CsCl (+CsCl, magenta) to fully block the channel. (**B**) Mean time courses of normalized current amplitudes of HCN1 wt 45 (black), A387S (blue empty) and wt/A387S (blue), recorded at -110 mV, before and after perfusion at time = 0 of 30 µM ivabradine (black line, +iva). (**C**) Percentage of mean maximum block by 30 µM ivabradine of HCN1 wt and mutant: 76.6 ± 6.1% (wt, black), 55.7 ± 10.6% (AS, empty blue) and 69.8 ± 10.6% (wt/AS, blue). There is no statistical difference between the three values (One-Way ANOVA). Values are mean of n ≥ 3 experiments ± SEM. (**D**) Representative current traces of HCN1 A387S channels in control solution (blue empty circle) or incubated with 60 µM Org-34167 in the extracellular solution (green circle). Wild type (wt, black circle) currents are shown for comparison to highlight the instantaneous component in A387S. Traces shown from -20 mV to -120 mV. Scale bars: 250 pA and 500 ms. (**E**) Corresponding mean I/V relationships of steady-state current density ($I_{SS}$, normalized) and (**F**) of instantaneous current ($I_{INST}$, normalized) of wt (black), A387S (blue empty) and A387S +60 µM Org-34167 channels showing that both current components are reduced by Org-34167. Data are mean ± SEM.

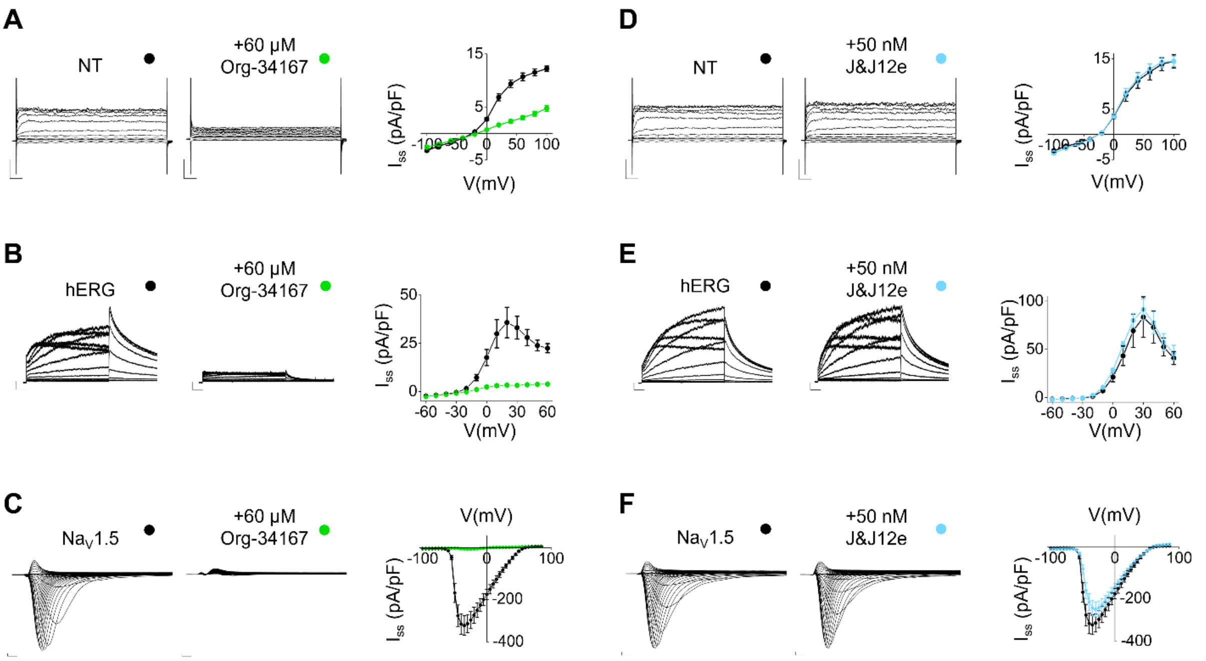


**Supplementary Figure 5. Org-34167, but not J&J12e, aspecifically blocks endogenous HEK293F currents, hERG, and Na$_V$1.5 channels.** Representative whole-cell currents and corresponding I/V relationships of untranfected (NT) HEK293F cells, HEK293F cells transfected with hERG and Na$_V$1.5 channels in control solution (black dot) or in presence of 60 µM Org-34167 (green dot) (**A-C**) or 50 nM J&J12e (light blue dot) (**D-F**). Traces shown from -100 mV to +100 mV for NT with scale bars: 200 pA and 25 ms. Data points are mean ± SEM. $I_{ss}$ values (in pA/pF) measured at +100 mV are as follows: wt = 12.2 ± 0.5 vs Org-34167 = 4.5 ± 0.6 (p = 1.6E-6) (**A**) and wt = 14.5 ± 1.0 vs J&J12e = 14.5 ± 1.3 (p = 1.0) (**D**). Traces shown from -60 mV to +60 mV for hERG channel with scale bars: 150 pA and 500 ms. Data points are mean ± SEM. $I_{ss}$ values (in pA/pF) measured at +20 mV are as follows: wt = 35.7 ± 7.9 vs Org-34167 = 3.3 ± 0.5 (p = 0.003) (**B**) and wt = 69.2 ± 17.4 vs J&J12e = 79.9 ± 7.0 (p = 0.5) (**E**). Traces shown from -100 mV to +85 mV for Na$_V$1.5 channel with scale bars: 250 pA and 0.5 ms. Data points are mean ± SEM. $I_{ss}$ values (in pA/pF) measured at -35 mV are as follows: wt = 329.5 ± 41.7 vs Org-34167 = -5.5 ± 1.8 (p = 6.6E-6) (**C**) and wt = 329.5 ± 41.7 vs J&J12e = -258.1 ± 33.3 (p = 0.3) (**F**). Statistical analysis was performed using a two-tailed Student's t-test.

#### Supplementary Tables

**Supplementary Table 1: Fitting parameters of the activation curve of HCN1 wt and mutant channels.** Half activation voltage ($V_{1/2}$) and inverse slope factor (k) values obtained by fitting data to a Boltzmann function (see Material and Methods) of HCN1 wt, HCN1 mutant (homotetramer) and HCN1 wt + HCN1 mutant (heterotetramer); n: number of cells tested in each condition. ND = not determined (HCN current absent or too small for analysis). NA = not analyzed. $\Delta V_{1/2}$: mutation-induced shift in $V_{1/2}$ compared to wild type, in mV. Each set of experiments contains data from controls and mutants measured on the same day. Values are reported as mean ± S.E.M. p values were calculated by Student's two-tailed unpaired T-test (§) or One Way ANOVA with Fisher's test (*) and compared to wt HCN1 in control condition. Statistically different p values ($\leq 0.05$) are highlighted in red.

| | Wild type | | | Homotetramer | | | | Heterotetramer | | | |
|---|---|---|---|---|---|---|---|---|---|---|---|
| | $V_{1/2}$ (mV) ± SEM | k (mV) ± SEM | n | $V_{1/2}$ (mV) ± SEM | k (mV) ± SEM | n | $\Delta V_{1/2}$ (mV) ± SEM | $V_{1/2}$ (mV) ± SEM | k (mV) ± SEM | n | $\Delta V_{1/2}$ (mV) ± SEM |
| **Q97R** | -71.6 ± 1.1 | 6.8 ± 0.4 | 7 | -78.8 ± 0.6 | 6.1 ± 0.6$^{\S p=0.3}$ | 6 | -7.2 ± 1.2$^{*p=6.6E-6}$ | -77.4 ± 0.6 | 6.3 ± 0.2$^{\S p=0.3}$ | 9 | -5.8 ± 1.1$^{*p=3.2E-5}$ |
| **S100A** | -72.1 ± 0.5 | 7.0 ± 1.2 | 4 | -70.5 ± 0.7 | 7.7 ± 1.3$^{\S p=0.6}$ | 5 | 1.6 ± 0.9$^{\S p=0.1}$ | NA | NA | | NA |
| **T172P** | -80.1 ± 1.3 | 6.2 ± 0.2 | 8 | ND | ND | | ND | -72.6 ± 1.9 | 7.6 ± 0.7$^{\S p=0.08}$ | 6 | +7.4 ± 2.3$^{\S p=0.007}$ |
| **L157V** | -71.5 ± 1.2 | 5.7 ± 0.4 | 6 | -70.8 ± 1.0 | 4.9 ± 0.3$^{\S p=0.2}$ | 6 | 0.7 ± 1.5$^{\S p=0.7}$ | -72.6 ± 1.0 | 5.5 ± 0.4$^{\S p=0.7}$ | 7 | -1.1 ± 1.5$^{\S p=0.5}$ |
| **W175R** | -77.3 ± 1.1 | 6.9 ± 0.6 | 10 | ND | ND | | ND | ND | ND | | ND |
| **N179Y** | -74.9 ± 1.0 | 7.6 ± 0.7 | 10 | ND | ND | | ND | -65.3 ± 0.7 | 7.6 ± 0.4$^{\S p=0.9}$ | 12 | +9.6 ± 1.2$^{\S p=8.5E-8}$ |
| **F186L** | -81.2 ± 0.9 | 7.8 ± 0.7 | 8 | -70.2 ± 2.8 | 6.6 ± 1.0$^{*p=0.3}$ | 5 | +11.0 ± 2.5$^{*p=3.7E-4}$ | -76.4 ± 1.6 | 8.1 ± 0.7$^{*p=0.8}$ | 8 | +4.8 ± 2.2$^{*p=0.04}$ |
| **D189Y** | -76.7 ± 0.6 | 6.8 ± 0.4 | 10 | ND | ND | | ND | -75.1 ± 0.7 | 6.1 ± 0.3$^{*p=0.2}$ | 10 | +1.6 ± 1.0$^{*p=0.1}$ |
| **N200S** | -79.2 ± 1.6 | 6.9 ± 0.5 | 10 | -78.2 ± 1.3 | 6.0 ± 0.4$^{*p=0.1}$ | 11 | 1.1 ± 2.1$^{*p=0.6}$ | -78.9 ± 1.4 | 6.4 ± 0.3$^{*p=0.0.4}$ | 13 | +0.4 ± 2.0$^{*p=0.8}$ |
| **I206V** | -74.4 ± 1.3 | 6.9 ± 0.6 | 14 | -79.8 ± 1.6 | 6.6 ± 0.4$^{*p=0.7}$ | 10 | -5.3 ± 2.1$^{*p=0.02}$ | -80.7 ± 1.7 | 7.0 ± 0.4$^{*p=0.9}$ | 9 | -6.3 ± 2.1$^{*p=0.007}$ |
| **M215V** | -74.4 ± 1.9 | 6.7 ± 0.5 | 5 | -75.0 ± 1.4 | 5.4 ± 0.3$^{\S p=0.1}$ | 4 | -0.2 ± 0.8$^{\S p=0.8}$ | NA | NA | | NA |
| **L237P** | -73.2 ± 1.0 | 6.2 ± 0.2 | 11 | ND | ND | | ND | -72.8 ± 0.6 | 6.9 ± 0.4$^{\S p=0.09}$ | 10 | -0.3 ± 1.2$^{\S p=0.8}$ |
| **E246K** | -79.8 ± 1.1 | 6.9 ± 0.3 | 10 | -71.8 ± 0.9 | 7.0 ± 0.3$^{*p=0.8}$ | 16 | +8.1 ± 1.4$^{*p=1.0E-6}$ | -69.4 ± 0.8 | 6.9 ± 0.3$^{*p=1.0}$ | 13 | +10.4 ± 1.4$^{*p=1.6E-8}$ |
| **K261E** | -74.7 ± 0.5 | 5.7 ± 0.3 | 6 | ND | ND | | ND | -75.2 ± 1.2 | 6.3 ± 0.6$^{*p=0.4}$ | 9 | -0.5 ± 1.5$^{\S p=0.7}$ |
| **A294V** | -78.4 ± 1.2 | 7.7 ± 0.3 | 9 | ND | ND | | ND | -66.5 ± 2.1 | 10.7 ± 0.8$^{\S p=0.01}$ | 7 | +11.9 ± 2.3$^{\S p=1.6E-4}$ |
| **S354N** | -74.32 ± 0.9 | 6.3 ± 0.3 | 18 | ND | ND | | ND | -85.1 ± 1.0 | 9.0 ± 0.5$^{\S p=4.2E-5}$ | 12 | -10.8 ± 1.4$^{\S p=3.6E-8}$ |

| | | | | | | | | | | | |
|---|---|---|---|---|---|---|---|---|---|---|---|
| **C358F** | -73.1 ± 1.5 | 6.3 ± 0.4 | 5 | ND | ND | | ND | -103.0 ± 1.3 | 9.9 ± 1.9 $^{\S p=0.006}$ | 5 | -29.9 ± 2.0 $^{\S p=4.3E-7}$ |
| **C358R** | -75.4 ± 0.7 | 7.5 ± 0.3 | 12 | ND | ND | | ND | -84.1 ± 1.3 | 7.1 ± 0.5 $^{\S p=0.5}$ | 13 | -8.7 ± 1.5 $^{\S p=6.5E-6}$ |
| **C358Y** | -75.5 ± 0.8 | 6.9 ± 0.5 | 11 | ND | ND | | ND | -76.0 ± 0.9 | 6.1 ± 0.7 $^{\S p=0.4}$ | 4 | +0.4 ± 1.5 $^{\S p=0.8}$ |
| **Y361C** | -75.6 ± 1.0 | 6.8 ± 0.5 | 12 | ND | ND | | ND | ND | ND | | ND |
| **M379R** | -73.7 ± 0.6 | 6.9 ± 0.2 | 8 | ND | ND | | ND | -72.9 ± 1.2 | 6.2 ± 0.8 $^{\S p=0.4}$ | 7 | -0.7 ± 1.3 $^{\S p=0.6}$ |
| **I380F** | -73.8 ± 0.8 | 6.8 ± .5 | 8 | ND | ND | | ND | -61.0 ± 1.8 | 10.6 ± 0.5 $^{\S p=3.3E-4}$ | 7 | -12.8 ± 1.9 $^{\S p=1.6E-5}$ |
| **A387S** | -79.5 ± 1.2 | 7.3 ± 0.3 | 8 | ND | ND | | ND | -63.1 ± 2.7 | 10.0 ± 1.3 $^{\S p=0.05}$ | 6 | +16.5 ± 2.7 $^{\S p=6.0E-5}$ |
| **A387G** | -72.1 ± 1.5 | 6.2 ± 0.4 | 7 | ND | ND | | ND | -61.5 ±1.6 | 9.5 ± 0.5 $^{\S p=1.9E-4}$ | 9 | +10.5 ± 2.2 $^{\S p=3.1E-4}$ |
| **Y411C** | -75.7 ± 1.1 | 7.1 ± 0.5 | 10 | -63.1 ± 0.8 | 6.9 ± 0.5 $^{*p=0.7}$ | 10 | +12.5 ± 1.3 $^{*p=8.9E-10}$ | -71.2 ± 0.8 | 6.5 ± 0.7 $^{*p=0.4}$ | 7 | +4.4 ± 1.4 $^{*p=0.004}$ |
| **D433Y** | -74.3 ± 0.5 | 6.2 ± 0.2 | 11 | ND | ND | | ND | -63.4 ± 1.7 | 8.3 ± 0.4 $^{\S p=6.0E-4}$ | 10 | +10.9 ± 1.7 $^{\S p=2.7E-6}$ |
| **L450P** | -70.6 ± 1.2 | 6.1 ± 0.2 | 7 | ND | ND | | ND | -71.9 ± 0.9 | 6.4 ± 0.3 $^{\S p=0.4}$ | 7 | -1.3 ± 1.5 $^{\S p=0.4}$ |
| **C542F** | -75.8 ± 0.6 | 6.1 ± 0.4 | 12 | -69.2 ± 1.4 | 11.3 ± 1.1 $^{*p=9.5E-6}$ | 8 | +6.6 ± 1.3 $^{*p=2.3E-5}$ | -72.8 ± 0.8 | 7.5 ± 0.5 $^{*p=0.1}$ | 8 | +3.0 ± 1.3 $^{*p=0.02}$ |
| **R548H** | -75.9 ± 0.8 | 6.6 ± 0.3 | 14 | -83.7 ± 1.2 | 7.5 ± 0.7 $^{*p=0.2}$ | 7 | -7.8 ± 1.4 $^{*p=1.2E-5}$ | -80.8 ± 1.3 | 6.7 ± 0.4 $^{*p=0.8}$ | 7 | -4.9 ± 1.4 $^{*p=0.002}$ |
| **R590Q** | -75.4 ± 1.0 | 7.7 ± 0.2 | 8 | -81.5 ± 1.0 | 7.1 ± 0.5 $^{*p=0.4}$ | 8 | -6.1 ± 1.5 $^{*p=6.7E-4}$ | -79.9 ± 1.4 | 7.0 ± 1.1 $^{*p=0.4}$ | 6 | -4.5 ± 1.6 $^{*p=0.01}$ |

**Supplementary Table 2: Mean steady state current density of HCN1 wt and mutant channels.** Mean steady state current density value measured after a -120 mV hyperpolarizing step normalized to each cell's capacitance ($I_{ss}$ at -120 mV, in pA/pF) of HCN1 wt, HCN1 mutant (homotetramer) and HCN1 wt + HCN1 mutant (heterotetramer); n: number of cells tested in each condition. NA = not analyzed. Each set of experiments contains data from controls and mutants measured on the same day. Values are reported as mean ± S.E.M. p values were calculated by Student's two-tailed unpaired T-test (§) or One Way ANOVA with Fisher's test (*) and compared to wt hHCN1 in control condition. Statistically different p values ($\leq 0.05$) are highlighted in red.

| | Wild type | | Homotetramer | | Heterotetramer | |
|---|---|---|---|---|---|---|
| | $I_{ss}$ (pA/pF) at -120 mV ± SEM | n | $I_{ss}$ (pA/pF) at -120 mV ± SEM | n | $I_{ss}$ (pA/pF) at -120 mV ± SEM | n |
| **Q97R** | -90.0 ± 14.8 | 6 | -91.2 ± 26.6 $^{*p=0.9}$ | 6 | -82.9 ± 17.4 $^{*p=0.8}$ | 6 |
| **S100A** | -118.8 ± 9.5 | 4 | -132.2 ± 11.7 $^{\S p=0.4}$ | 4 | NA | |
| **L157V** | -102.3 ± 20.1 | 8 | -42.3 ± 12.3 $^{*p=0.01}$ | 7 | -54.4 ± 11.5 $^{*p=0.03}$ | 8 |
| **T172P** | -135.5 ± 10.2 | 10 | -30.7 ± 7.8 $^{*p=1.3E-4}$ | 4 | -57.4 ± 13.9 $^{*p=1.4E-4}$ | 11 |
| **W175R** | -133.0 ± 31.3 | 10 | -4.9 ± 0.6 $^{*p=0.004}$ | 5 | -4.5 ± 0.3 $^{*p=0.005}$ | 6 |
| **N179Y** | -131.1 ± 28.5 | 10 | -15.7 ± 1.2 $^{*p=3.8E-4}$ | 17 | -48.9 ± 16.5 $^{*p=0.005}$ | 8 |
| **F186L** | -249.2 ± 57.8 | 9 | -107.6 ± 33.8 $^{*p=0.08}$ | 5 | -261.4 ± 40.6 $^{*p=0.8}$ | 8 |
| **D189Y** | -116.3 ± 18.8 | 10 | -6.3 ± 0.8 $^{*p=9.9E-8}$ | 12 | -63.3 ± 8.8 $^{*p=0.003}$ | 10 |
| **N200S** | -172.7 ± 32.1 | 10 | -155.7 ± 45.1 $^{*p=0.7}$ | 11 | -160.4 ± 21.2 $^{*p=0.8}$ | 13 |
| **I206V** | -130 ± 19.1 | 14 | -121.8 ± 29.9 $^{*p=0.8}$ | 10 | -128.8 ± 32.6 $^{*p=0.9}$ | 9 |
| **M215V** | -65.5 ± 18.9 | 5 | -58.8 ± 7.4 $^{\S p=0.7}$ | 5 | NA | |
| **L237P** | -87.5 ± 17.9 | 10 | -8.6 ± 1.4 $^{*p=1.5E-4}$ | 11 | -90.0 ± 14.3 $^{*p=0.9}$ | 10 |
| **E246K** | -169.5 ± 32.9 | 10 | -173.6 ± 28.6 $^{*p=0.9}$ | 15 | -169.1 ± 39.4 $^{*p=1.0}$ | 13 |
| **K261E** | -105.9 ± 21.2 | 6 | -5.9 ± 0.7 $^{*p=0.007}$ | 8 | -114.1 ± 32.4 $^{*p=0.8}$ | 8 |
| **A294V** | -263.7 ± 62.9 | 10 | -206.4 ± 24.3 $^{*p=0.2}$ | 8 | -182.0 ± 43.4 $^{*p=0.4}$ | 12 |
| **S354N** | -91.9 ± 19.7 | 19 | -16.1 ± 2.8 $^{*p=0.003}$ | 11 | -89.3 ± 12.8 $^{*p=0.9}$ | 17 |
| **C358F** | -116.7 ± 24.0 | 19 | -19.3 ± 2.8 $^{*p=0.005}$ | 16 | -49.7 ± 9.8 $^{*p=0.03}$ | 9 |
| **C358R** | -159.1 ± 14.6 | 18 | -14.1 ± 3.7 $^{*p=5.2E-9}$ | 15 | -82.9 ± 15.2 $^{*p=2.4E-4}$ | 15 |
| **C358Y** | -111.0 ± 22.6 | 17 | -5.5 ± 0.6 $^{*p=1.3E-4}$ | 11 | -28.4 ± 7.5 $^{*p=0.002}$ | 11 |
| **Y361C** | -131.9 ± 21.5 | 17 | -8.0 ± 1.6 $^{*p=4.0E-5}$ | 12 | -9.5 ± 2.1 $^{*p=2.5E-4}$ | 6 |

| | | | | | | |
|---|---|---|---|---|---|---|
| **M379R** | -152.9 ± 26.3 | 8 | -6.4 ± 1.0 *p=9.1E-6 | 8 | -38.1 ± 16.0 *p=2.4E-4 | 7 |
| **I380F** | -116.3 ± 20.2 | 7 | -110.5 ± 17.2 *p=0.8 | 12 | -88.5 ± 18.5 *p=0.3 | 8 |
| **A387S** | -169.9 ± 37.5 | 9 | -140.6 ± 19.8 *p=0.5 | 14 | -167.4 ± 4.1 *p=1.0 | 10 |
| **A387G** | -152.2 ± 29.2 | 7 | -64.2 ± 10.9 *p=0.003 | 8 | -84.2 ± 13.5 *p=0.02 | 8 |
| **Y411C** | -141.6 ± 35.9 | 10 | -154.5 ± 59.0 *p=0.8 | 9 | -146.5 ± 48.1 *p=0.9 | 7 |
| **D433Y** | -94.9 ± 12.3 | 10 | -59.9 ± 9.2 *p=0.03 | 15 | -58.1 ± 13.1 *p=0.04 | 9 |
| **L450P** | -112.2 ± 7.2 | 6 | -31.4 ± 14.4 *p=0.008 | 6 | -115.1 ± 21.4 *p=0.9 | 7 |
| **C542F** | -91.4 ± 16.0 | 12 | -100.0 ± 22.8 *p=0.8 | 8 | -89.1 ± 22.7 *p=0.9 | 9 |
| **R548H** | -103.2 ± 15.7 | 11 | -106.9 ± 23.7 *p=0.9 | 7 | -97.7 ± 21.3 *p=0.8 | 6 |
| **R590Q** | -448.7 ± 116.4 | 8 | -200.2 ± 51.7 *p=0.04 | 9 | -167.4 ± 37.5 *p=0.04 | 5 |

**Supplementary Table 3: Co-localization parameters of wild type and mutant HCN1.** Summary of mean Pearson correlation coefficients (r) and peak distance (PD) values of wild-type and mutant HCN1 channels. Mean r and PD were calculated on the intensity profiles of fluorescent signals encompassing the plasma membrane obtained as described in Material and Methods. n: number of cells. Each set of experiments contains data from controls and mutants imaged on the same day. Values are reported as mean ± S.E.M.

| | **Homotetramer** | | | **Heterotetramer** | | |
|---|---|---|---|---|---|---|
| | **r ± SEM** | **PD ± SEM (μm)** | **n** | **r ± SEM** | **PD ± SEM (μm)** | **n** |
| **wild-type** | 0.32 ± 0.02 | 0.03 ± 0.01 | 5 | | | 5 |
| **W175R** | 0.25 ± 0.1 | 1.02 ± 0.4 | 5 | 0.09 ± 0.2 | 1.0 ± 0.2 | 5 |
| **N179Y** | 0.37 ± 0.05 | 0.46 ± 0.08 | 5 | 0.67 ± 0.07 | 0.02 ± 0.02 | 5 |
| **D189Y** | 0.36 ± 0.07 | 1.0 ± 0.4 | 5 | 0.78 ± 0.02 | 0.12 ± 0.05 | 5 |
| **I206V** | 0.62 ± 0.07 | 0.15 ± 0.04 | 5 | 0.88 ± 0.03 | 0.11 ± 0.07 | 5 |
| **L237P** | 0.39 ± 0.13 | 0.77 ± 0.3 | 5 | 0.70 ± 0.05 | 0.55 ± 0.41 | 5 |
| **K261E** | 0.30 ± 0.04 | 0.99 ± 0.1 | 5 | 0.77 ± 0.02 | 0.18 ± 0.06 | 5 |
| **S354N** | 0.80 ± 0.02 | 0.11 ± 0.03 | 5 | 0.94 ± 0.01 | 0.11 ± 0.05 | 5 |
| **C358F** | 0.79 ± 0.02 | 0.06 ± 0.02 | 5 | 0.91 ± 0.01 | 0.11 ± 0.04 | 5 |
| **C358R** | 0.86 ± 0.01 | 0.07 ± 0.03 | 5 | 0.92 ± 0.03 | 0.07 ± 0.04 | 5 |
| **C358Y** | 0.88 ± 0.03 | 0.10 ± 0.03 | 5 | 0.83 ± 0.03 | 0.03 ± 0.001 | 5 |
| **Y361C** | 0.88 ± 0.03 | 0.09 ± 0.02 | 5 | 0.89 ± 0.03 | 0.06 ± 0.02 | 5 |
| **M379R** | 0.38 ± 0.17 | 0.58 ± 0.2 | 5 | 0.78 ± 0.04 | 0.27 ± 0.01 | 5 |
| **A387G** | 0.58 ± 0.12 | 0.22 ± 0.09 | 5 | 0.90 ± 0.04 | 0.13 ± 0.07 | 5 |
| **D433Y** | 0.97 ± 0.007 | 0.05 ± 0.02 | 5 | 0.90 ± 0.02 | 0.09 ± 0.02 | 5 |
| **L450P** | 0.46 ± 0.09 | 0.40 ± 0.17 | 5 | 0.83 ± 0.07 | 0.08 ± 0.07 | 5 |

**Supplementary Table 4: Summary of $F_{PR}/F_{PM}$ ratio values of wild type and mutant HCN1.** Mean $F_{PR}/F_{PM}$ ratio of the fluorescence signal of eGFP-HCN1 wt or mutant measured at the perinuclear ring (PR) and at the plasma membrane (PM). $F_{PR}/F_{PM}$ ratio for each cell calculated as the mean ratio of 10 points on the PR and PM, respectively. n: number of cells tested in each condition. Each set of experiments contains data from controls and mutants imaged on the same day. Values are reported as mean ± S.E.M. *p values were calculated by One Way ANOVA with Fisher's test and compared to wt HCN1. Statistically different p values (≤ 0.05) are highlighted in red.

| | Wild type | | Homotetramer | | Heterotetramer | |
|---|---|---|---|---|---|---|
| | $F_{PR}/F_{PM}$ ± SEM | n | $F_{PR}/F_{PM}$ ± SEM | n | $F_{PR}/F_{PM}$ ± SEM | n |
| **W175R** | 0.5 ± 0.03 | 5 | 2.7 ± 0.03 *p=1.9E-13 | 5 | 2.0 ± 0.07 *p=1.7E-11 | 10 |
| **N179Y** | 0.5 ± 0.03 | 5 | 1.0 ± 0.02 *p=2.5E-9 | 5 | 1.0 ± 0.01 *p=6.7E-9 | 5 |
| **D189Y** | 0.5 ± 0.01 | 5 | 3.0 ± 0.07 *p=3.6E-14 | 5 | 0.7 ± 0.003 *p=0.003 | 5 |
| **I206V** | 0.5 ± 0.03 | 5 | 0.7 ± 0.06 *p=0.03 | 5 | 0.5 ± 0.01 *p=0.5 | 5 |
| **L237P** | 0.5 ± 0.008 | 5 | 2.4 ± 0.05 *p=3.2E-14 | 5 | 0.7 ± 0.05 *p=3.8E-4 | 5 |
| **K261E** | 0.5 ± 0.01 | 5 | 3.3 ± 0.1 *p=8.1E-12 | 5 | 1.1 ± 0.03 *p=8.8E-5 | 5 |
| **S354N** | 0.5 ± 0.03 | 5 | 0.6 ± 0.03 *p=0.6 | 5 | 0.5 ± 0.01 *p=0.9 | 5 |
| **C358F** | 0.5 ± 0.03 | 5 | 0.5 ± 0.05 *p=0.4 | 5 | 0.5 ± 0.06 *p=0.1 | 5 |
| **C358R** | 0.5 ± 0.03 | 5 | 0.7 ± 0.06 *p=0.01 | 5 | 0.7 ± 0.04 *p=0.005 | 5 |
| **C358Y** | 0.4 ± 0.03 | 5 | 0.5 ± 0.03 *p=0.3 | 5 | 0.4 ± 0.05 *p=0.5 | 5 |
| **Y361C** | 0.5 ± 0.03 | 5 | 0.9 ± 0.04 *p=2.9E-6 | 5 | 0.6 ± 0.04 *p=4.4E-7 | 5 |
| **M379R** | 0.5 ± 0.02 | 5 | 2.6 ± 0.3 *p=7.3E-7 | 5 | 0.6 ± 0.04 *p=0.6 | 5 |
| **A387G** | 0.4 ± 0.01 | 5 | 0.5 ± 0.03 *p=0.01 | 5 | 0.5 ± 0.01 *p=0.02 | 5 |
| **D433Y** | 0.4 ± 0.04 | 5 | 0.5 ± 0.04 *p=0.08 | 5 | 0.4 ± 0.03 *p=0.9 | 5 |
| **L450P** | 0.5 ± 0.03 | 5 | 1.1 ± 0.1 *p=7.9E-4 | 5 | 1.0 ± 0.1 *p=0.004 | 5 |

**Supplementary Table 5: Fitting parameters of the activation curves of HCN1 wt and mutant channels treated with NB6 and TRIP8b$_{nano}$.** Half activation voltage ($V_{1/2}$) and inverse slope factor (k) obtained by fitting data to a Boltzmann function (see Material and Methods) in absence (control) or in presence of 20 µM NB6 or 10 µM TRIP8b$_{nano}$. Treatment: 20 µM NB6 added to the extracellular recording solution, or 10 µM TRIP8b$_{nano}$ added in the pipette solution, in nanomolar (nM); n: number of cells tested in each condition. $\Delta V_{1/2}$: treatment-induced shift in $V_{1/2}$, in mV, compared to wild-type channels. Each set of experiments contains data from controls and J&J12e measured on the same day. Values are reported as mean ± S.E.M. *$p<0.05$ values were calculated by One-way ANOVA with Fisher's test compared to hHCN1 wt or mutant in control solution; §p values were calculated by Student's two-tailed unpaired T-test compared to control condition. Statistically different p values (≤ 0.05) are highlighted in red.

| | Wild type | | | Heterotetramer | | | | +Treatment | | | |
|---|---|---|---|---|---|---|---|---|---|---|---|
| | **$V_{1/2}$ (mV) ± SEM** | **k (mV) ± SEM** | **n** | **$V_{1/2}$ (mV) ± SEM** | **k (mV) ± SEM** | **n** | **$\Delta V_{1/2}$ (mV) ± SEM** | **$V_{1/2}$ (mV) ± SEM** | **k (mV) ± SEM** | **n** | **$\Delta V_{1/2}$ (mV) ± SEM** |
| **wt +NB6** | -73.2 ± 1.2 | 6.7 ± 0.3 | 5 | | | | | -65.8 ± 1.3 | 6.3 ± 0.3$^{§p=0.7}$ | 4 | 7.4 ± 1.8$^{§p=0.004}$ |
| **wt/R548H +NB6** | -73.2 ± 0.6 | 6.1 ± 0.4 | 7 | -78.0 ± 0.4 | 6.3 ± 0.6$^{*p=0.7}$ | 7 | -4.6 ± 0.8$^{*p=1.0E-5}$ | -71.3 ± 0.6 | 6.0 ± 0.4$^{*p=0.9}$ | 7 | 2.0 ± 0.8$^{*p=0.06}$ |
| **wt/S354N +NB6** | -72.2 ± 0.8 | 6.2 ± 0.3 | 11 | -84.0 ± 1.3 | 9.4 ± 0.7$^{*p=8.6E-4}$ | 8 | -11.8 ± 1.5$^{*p=1.8E-7}$ | -74.9 ± 1.8 | 10.1 ± 1.2$^{*p=5.5E-4}$ | 5 | -2.7 ± 1.8$^{*p=0.1}$ |
| **wt + TRIP8b$_{nano}$** | -72.4 ± 0.7 | 7.7 ± 0.9 | 8 | | | | | -80.3 ± 0.9 | 7.5 ± 0.5$^{*p=0.8}$ | 7 | -7.8 ± 1.2$^{§p=1.5E-5}$ |
| **wt/E246K + TRIP8b$_{nano}$** | -73.1 ± 0.9 | 6.3 ± 0.5 | 6 | -65.0 ± 1.0 | 7.0 ± 0.6$^{*p=0.3}$ | 6 | 8.1 ± 1.2$^{*p=5.4E-6}$ | -71.3 ± 0.6 | 6.9 ± 0.4$^{*p=0.3}$ | 7 | 1.7 ± 1.2$^{*p=0.1}$ |
| **wt/M153I + TRIP8b$_{nano}$** | -72.4 ± 0.7 | 7.7 ± 0.9 | 8 | -67.5 ± 0.8 | 8.3 ± 0.2$^{*p=0.6}$ | 8 | 14.9 ± 1.0$^{*p=1.5E-12}$ | -65.0 ± 0.6 | 8.6 ± 0.5$^{*p=0.3}$ | 8 | 7.4 ± 1.0$^{*p=3.1E-7}$ |

**Supplementary Table 6: Mean steady state current density of HCN1 wt and mutant channels treated with NB6 and TRIP8b$_{nano}$.** Mean steady state current density value measured after a -120 mV hyperpolarizing step normalized to each cell's capacitance ($I_{ss}$ at -120 mV, in pA/pF) of HCN1 wt and HCN1 wt + HCN1 mutant (heterotetramer) in absence (control) or in presence of 20 µM NB6 or 10 µM TRIP8b$_{nano}$. Treatment: 20 µM NB6 added to the extracellular recording solution, or 10 µM TRIP8b$_{nano}$ added in the pipette solution, in nanomolar (nM). n: number of cells tested in each condition. Each set of experiments contains data from controls and mutants measured on the same day. Values are reported as mean ± S.E.M. p values were calculated by Student's two-tailed unpaired T-test (§) or One Way ANOVA with Fisher's test (*) and compared to wt hHCN1 in control condition. Statistically different p values (≤ 0.05) are highlighted in red.

| | Wild type | | Heterotetramer | | +Treatment | |
|---|---|---|---|---|---|---|
| | $I_{SS}$ (pA/pF) at -120 mV ± SEM | n | $I_{SS}$ (pA/pF) at -120 mV ± SEM | n | $I_{SS}$ (pA/pF) at -120 mV ± SEM | n |
| **wt +NB6** | -136.2 ± 41.8 | 5 | | | -142.4 ± 33.8 $^{§p=0.9}$ | 5 |
| **wt/R548H +NB6** | -94.6 ± 27.6 | 6 | -98.6 ± 22.0 $^{*p=0.9}$ | 7 | -93.8 ± 12.5 $^{*p=1.0}$ | 8 |
| **wt/S354N +NB6** | -68.8 ± 10.8 | 11 | -67.7 ± 13.1 $^{*p=0.9}$ | 7 | -68.0 ± 11.6 $^{*p=0.9}$ | 5 |
| **wt + TRIP8b$_{nano}$** | -94.8 ± 16.3 | 8 | | | -94.0 ± 12.7 $^{*p=1.0}$ | 7 |
| **wt/E246K + TRIP8b$_{nano}$** | -108.5 ± 23.3 | 6 | -112.4 ± 33.2 $^{*p=0.9}$ | 6 | -111.7 ± 23.5 $^{*p=0.9}$ | 7 |
| **wt/M153I + TRIP8b$_{nano}$** | -94.8 ± 16.3 | 8 | -91.0 ± 14.0 $^{*p=0.9}$ | 9 | -94.8 ± 17.3 $^{*p=1.0}$ | 8 |

**Supplementary Table 7: Fitting parameters of the activation curves and I/V relationships of HCN1 wt and HCN1 wt / M153I channels treated with J&J12e.** Half activation voltage ($V_{1/2}$), inverse slope factor (k) and maximal tail amplitude values (at -120 mV) obtained by fitting data to a Boltzmann function (see Material and Methods) and mean steady state current density value measured after a -130 mV hyperpolarizing step normalized to each cell's capacitance ($I_{ss}$ at -130 mV, in pA/pF) in absence (control) or in presence of J&J12e (12e). $I_{ss}$ values are normalized on a 0-100 scale, hHCN1 wt and mutant channels in control solution are assigned a value of $I_{ss}$ (pA/pF) at -130 mV = 100. [J&J12e]: concentration of J&J12e added to the extracellular recording solution, in nanomolar (nM); n: number of cells tested in each condition. $\Delta V_{1/2}$: J&J12e-induced shift in $V_{1/2}$, in mV. Each set of experiments contains data from controls and J&J12e measured on the same day. Values are reported as mean ± S.E.M. *$p<0.05$ values were calculated by One-way ANOVA with Fisher's test compared to hHCN1 wt or mutant in control solution; §p values were calculated by Student's two-tailed unpaired T-test compared to control condition. Statistically different p values (≤ 0.05) are highlighted in red.

| Construct | $V_{1/2}$ (mV) ± SEM control | k (mV) ± SEM control | n | [12e] (nM) | $V_{1/2}$ (mV) ± SEM with 12e | k (mV) ± SEM with 12e | n | $\Delta V_{1/2}$ (mV) ± SEM | Max tail ampl. (mV) ± SEM | $I_{ss}$ (pA/pF) at -130 mV ± SEM | n |
|---|---|---|---|---|---|---|---|---|---|---|---|
| hHCN1 wt | -75.9 ± 0.8 | 6.3 ± 0.2 | 16 | **1000** | ND | ND | | ND | ND | 5.1 ± 3.2$^{*p=0.01}$ | 5 |
| | | | | **100** | ND | ND | | ND | ND | 19.9 ± 8.5$^{*p=0.01}$ | 3 |
| | | | | **50** | -91.0 ± 3.2 | 8.8 ± 0.7 $^{*p=5.3E-4}$ | 4 | -15.2 ± 2.0 $^{*p=1.7E-8}$ | 3.5 ± 0.5 $^{*p=0.06}$ | 48.6 ± 19.5$^{*p=0.1}$ | 4 |
| | | | | **25** | -88.0 ± 0.4 | 8.7 ± 0.3 $^{*p=3.9E-4}$ | 5 | -12.1 ± 1.8 $^{*p=1.7E-8}$ | 4.4 ± 0.7 $^{*p=0.5}$ | 63.4 ± 23.6$^{*p=0.1}$ | 5 |
| | | | | **10** | -80.9 ± 1.3 | 7.1 ± 0.6 $^{*p=0.1}$ | 7 | -5.0 ± 1.6 $^{*p=0.004}$ | 6.3 ± 0.9 $^{*p=0.8}$ | 74.8 ± 13.1$^{*p=0.2}$ | 7 |
| | | | | **1** | -76.5 ± 1.6 | 7.4 ± 0.7 $^{*p=0.09}$ | 4 | -0.6 ± 2.0 $^{*p=0.07}$ | 6.5 ± 1.2 $^{*p=0.7}$ | 100.7 ± 8.1$^{*p=0.8}$ | 5 |
| hHCN1 wt / M153I | -56.6 ± 0.6 | 8.1 ± 0.3 | 12 | **1000** | ND | ND | | ND | ND | 13.6 ± 7.0$^{*p=0.02}$ | 4 |
| | | | | **100** | ND | ND | | ND | ND | 23.1 ± 7.9$^{*p=0.04}$ | 4 |
| | | | | **50** | -76.6 ± 1.6 | 9.4 ± 0.5 $^{*p=0.07}$ | 5 | -20.0 ± 1.5$^{*p=1.8E-14}$ | 5.0 ± 1.1 $^{*p=0.6}$ | 50.0 ± 12.2$^{*p=0.3}$ | 5 |
| | | | | **25** | -71.8 ± 1.2 | 7.7 ± 0.6 $^{*p=0.6}$ | 5 | -15.2 ± 1.5$^{*p=1.8E-11}$ | 5.8 ± 1.1 $^{*p=0.6}$ | 59.9 ± 20.1$^{*p=0.5}$ | 5 |
| | | | | **10** | -63.7 ± 1.3 | 7.8 ± 1.1 $^{*p=0.6}$ | 4 | -7.0 ± 1.6 $^{*p=1.2E-4}$ | 5.9 ± 2.9 $^{*p=0.6}$ | 78.7 ± 24.3$^{*p=0.2}$ | 5 |
| | | | | **1** | -57.0 ± 1.3 | 8.0 ± 0.3 $^{*p=0.9}$ | 3 | -0.4 ± 1.8 $^{*p=0.8}$ | 3.2 ± 0.5 $^{*p=0.9}$ | 99.5 ± 7.5$^{*p=0.6}$ | 4 |

**Supplementary Table 8: Functional classes and phenotypic severity distribution among probands with epilepsy (n = 41).** Epilepsy severity was categorized a priori as mild (generalized genetic epilepsy [GGE], febrile seizures plus [FS+], GEFS+, and focal epilepsies without encephalopathy), moderate (unclassified epilepsies, focal hemiclonic seizures, or a single episode of refractory status epilepticus), or severe (neonatal- or infantile-onset developmental and epileptic encephalopathy [DEE]). See Supplementary Methods for additional details. Percentages are calculated within each functional subgroup.

| Functional class | Mild n (%) | Moderate n (%) | Severe n (%) | Total (n) |
|---|---|---|---|---|
| LOF | 5 (62.5%) | 3 (37.5%) | 0 | 8 |
| Non-LOF | 4 (12.1%) | 4 (12.1%) | 25 (75.8%) | 33 |
| a) GOF | 2 (8.3%) | 3 (12.5 %) | 19 (79.2%) | 24 |
| b) LOF/GOF | 1 (16.7%) | 0 | 5 (83.3%) | 6 |
| c) wt-like | 1 (33.3%) | 1 (33.3%) | 1 (33.3%) | 3 |

LOF = loss-of-function, GOF = gain-of-function, DEE = developmental and epileptic encephalopathy, GGE = generalized genetic epilepsy, FS+ = febrile seizures plus, GEFS+ = genetic epilepsy with febrile seizures plus, SE = status epilepticus, n = number, wt = wild type.

#### Supplementary Table 9: Clinical summary of patients carrying HCN1 variants included in the study

| Functional Effect | PT-ID | Novel/ Ref | AA change | Age at time of study | Gender | Variant Segregation | Epilepsy (YES/NO) | Epilepsy/ syndrome type ILAE classification | Degree of severity of the whole phenotype epilepsy and DD | Age at seizure onset | Developmental delay (mild/moderate/ severe) | Other neuropsychiatric issues (ASD, ADHD, behavioural problems) | Other clinical features | MRI |
|---|---|---|---|---|---|---|---|---|---|---|---|---|---|---|
| LOF | 1 | novel | p.Gln97Arg | 9 yrs | M | maternally inherited (with sz) | YES | GGE with photosensitivity | mild | 9 yrs | mild | No | Bilateral hearing loss in 2023 | normal |
| wt-like | 2 | Kang et al. | p.Ser100Ala | 24 yrs | M | de novo | YES | Focal drug responsive epilepsy | mild | 15 yrs | NA | NA | NR | normal |
| GOF | 3 | Nava et al. Porro et al. | p.Ser100Phe | 6 yrs | F | de novo | YES | Infantile DEE (DS-like) | severe | 10 m | moderate | ASD | NR | normal |
| GOF /LOF | 4 | Marini et al. Porro et al. | p.Phe143Tyr | 12 yrs 6 m | F | de novo | YES | Infantile DEE | severe | 12 m | severe | NA | NR | normal |
| GOF | 5 | Marini et al. Porro et al. | p.Met153Ile | 7 yrs 3 m | F | de novo | YES | Infantile DEE | severe | 5 m | moderate | No | NR | normal |
| GOF | 6 | Marini et al. Porro et al. | p.Met153Ile | 2 yrs 8 m | M | de novo | YES | Unclassified infantile epilepsy | moderate | 9 m | No | language delay | NR | normal |
| LOF | 7 | Bonzanni et al. | p.Leu157Val | adult | M | de novo | YES | GGE | mild | 2 yrs (FS); 19 yrs | No | NA | NR | normal |
| GOF /LOF | 8 | Marini et al. | p.Thr172Pro | 42 yrs | F | de novo | YES | GGE-TCS | mild | 7 m | mild | ASD, behavioural problems | NR | normal |

| | | | | | | | | | | | | | | |
|---|---|---|---|---|---|---|---|---|---|---|---|---|---|---|
| LOF | 9 | novel | p.Asp189Tyr | 7 yrs | M | paternally inherited (with sz) | YES | FS+ | mild | 12 m | NA | No | NR | normal |
| wt-like§ | 10 | novel | p.Met215Val; pGly73_Gly74del | 10.5 yrs | F | p.Met215Val (paternal inheritance; pGly73_Gly74 del (maternally inherited compound heterozygous) | YES | Infantile DEE | severe | 10 m | severe | NA | hypotonia | normal |
| wt-like (LOF in homo) | 11 | novel | p.Leu237Pro | 4 yrs | M | maternally inherited (with learning problems) | YES | Unclassified infantile epilepsy | moderate | 12 m | moderate | No | NR | normal |
| wt-like (LOF in homo) | 12 | Marini et al. | p.Lys261Glu; c.1377+1G>A | 16 yrs | F | patient adopted | YES | Infantile DEE | severe | 7 m | severe | NA | wheelchair bound, cannot walk, hypotonia | Progressive cortical and white matter volume loss involving bilateral parietal/occipital lobes |
| GOF | 13 | Nava et al. Porro et al. | p.Ser272Pro | 16 yrs | F | de novo | YES | Infantile DEE (DS-like) | severe | 8 m | severe | ASD | behavioural disturbances | normal |
| GOF | 14 | novel | p.Ser272Pro | 8 yrs | M | de novo | YES | Focal epilepsy with hemiclonic prolonged seizures | severe | Infancy | DD | ASD, aggression & other neurobehavioral issues | pt underwent temporal lobectomy (tissue= gliosis and neuron loss) | subtle loss of gray-white matter differentiation within the anteromedial left temporal lobe, along with hippocampal volume loss and mild T2/FLAIR signal abnormality. Suggestive of anterior temporal cortical |

| | | | | | | | | | | | | | | |
|---|---|---|---|---|---|---|---|---|---|---|---|---|---|---|
| | | | | | | | | | | | | | | dysplasia and mesial temporal sclerosis, respectively. |
| GOF | 15 | Nava et al. Porro et al. | p.His279Tyr | 12 yrs | M | de novo | YES | Infantile DEE (DS-like) | severe | 13 m | mild | ADHD, behavioural disturbances, ataxia | NR | normal |
| GOF | 16 | novel | p.Ala294Val | 7 yrs | F | de novo | YES | Infantile DEE (DS-like) | severe | 5 m | moderate | DSA | NR | normal |
| GOF | 17 | Nava et al. Porro et al. | p.Arg297Thr | 15 yrs | F | de novo | YES | Infantile DEE (DS-like) | severe | 8 m | moderate>severe | ASD, behavioural problems | NR | normal |
| GOF | 18 | Marini et al. | p.Met305Leu | 1 yr | F | de novo | YES | Unclassfied early infantile epilepsy | moderate | 3 m | mild | NA | NR | normal |
| GOF | 19 | Marini et al. | p.Met305Leu | 14 yrs | F | de novo | YES | Infantile DEE | severe | 2 m | severe | NA | microcepahly | normal |
| LOF | 20 | novel | p.Ser354Asn | 3 yrs | F | de novo | NO | | | | mild | No | NR | normal |
| LOF | 21 | novel | p.Cys358Arg | 6 yrs | M | de novo | YES | Refractory SE (single episode) | moderate | | moderate | ASD | NR | normal |
| LOF | 22 | novel | p.Cys358Arg | 16 yrs | M | NA | YES | Early onset absence epilepsy | mild | | NA | behavioural problems | NR | normal |
| LOF | 23 | novel | p.Cys358Arg | 7 yrs | F | NA | NO | | | | moderate | No | hypotonia, dysmorphic features; mild ataxic gait | normal |

| | | | | | | | | | | | | | | |
|---|---|---|---|---|---|---|---|---|---|---|---|---|---|---|
| LOF | 24 | novel | p.Cys358Phe | 12 yrs 5 m | F | unknown (not inherited from the mother, father not tested) | NO | | | | severe | NA | bucofacial hypotonia, drooling | T2 hypersignals in bilateral occipital subcortical white matter |
| LOF | 25 | novel | p.Cys358Tyr | NA | F | de novo | NO | | | | moderate | ASD | NR | normal |
| LOF | 26 | novel | p.Tyr361Cys | 7 yrs | F | de novo | NO | | | | severe | ASD | NR | normal |
| LOF | 27 | Marini et al. | p.Met379Arg | NA | F | de novo | NO | | | | moderate | NA | NR | normal |
| GOF | 28 | Xie et al. | p.Ile380Phe | diet at 2 yrs | F | de novo | YES | Neonatal DEE | severe | 2 d | severe | NA | NR | normal |
| GOF | 29 | Wang et al. | p.Ile380Phe | NA | F | de novo | YES | Infantile DEE | severe | 50 days | severe | NA | NR | normal |
| GOF | 30 | novel | p.Ile380Phe | NA | F | de novo | YES | Infantile DEE | severe | 3 m | severe | No | NR | normal |
| GOF | 31 | novel | p.Ala387Ser | 6 yrs 5 m | M | unknown | YES | Infantile DEE | severe | 3m | severe | No | NR | normal |
| GOF | 32 | Lucariello et al. | p.Ala387Ser | 24 yrs | F | de novo | YES | Early infantile DEE | severe | 3 m | severe | NA | RETT like phenotype; microcephaly | NA |
| GOF | 33 | McKenzie et al. | p.Ala387Ser | 7 yrs | F | de novo | YES | Infantile DEE | severe | 4 m | severe | NA | NR | NA |

| | | | | | | | | | | | | | | |
|---|---|---|---|---|---|---|---|---|---|---|---|---|---|---|
| GOF /LOF | 34 | novel | p.Ala387Gly | 5 yrs | F | de novo | YES | infantile onset DEE | severe | 6 m | severe | No | NR | normal |
| GOF | 35 | novel | p.Gly391Ser | 32 yrs | F | maternally inherited (with sz) | YES | Infantile DEE | severe | NA | moderate | ASD | mild diplegy, constipation, shoulder luxations, obesity, dysmorphisms | mild broadening of the lateral ventricles |
| GOF | 36 | Marini et al. | p.Gly391Ser | 6 yrs 3 m | F | de novo | YES | FS+ | mild | 5 m | No | NA | NR | normal |
| GOF | 37 | Marini et al. | p.Gly391Ser | 2 yrs 5 m | M | de novo | YES | GEFS+ | mild | 7 m | mild | NA | NR | diffuse brain atrophy |
| GOF /LOF | 37 | Marini et al. | p.Gly391Asp | died at 14 m | M | de novo | YES | Neonatal DEE | severe | 30 h | severe | NA | NR | diffuse brain atrophy |
| GOF /LOF | 39 | Marini et al Fernández-Marmiesse A, et al. | p.Gly391Asp | died at 15 m | M | de novo | YES | Neonatal DEE | severe | 48 h | severe | NA | NR | diffuse brain atrophy |
| LOF | 40 | Marini et al. | p.Gly391Cys | 29 yrs | F | de novo | YES | GGE with eyelid myoclonia | moderate | Infancy | moderate | ASD | NR | diffuse brain atrophy |
| GOF | 41 | Marini et al. | p.Ile397Leu | 15 yrs | M | de novo | YES | Infantile DEE | severe | 5 m | severe | ASD | NR | diffuse brain atrophy |
| GOF | 42 | Marini et al. Porro et al. | p.Ser399Pro | 7 yrs | M | de novo | YES | Infantile DEE | severe | 4 m | severe | NA | NR | diffuse brain atrophy |
| GOF | 43 | Nava et al. Porro et al. | p.Asp401His | 18 yrs | F | de novo | YES | Infantile DEE (DS-like) | severe | 4 m | moderate to severe | ASD, behavioural problems | Polyphagia | normal |

| | | | | | | | | | | | | | |
|---|---|---|---|---|---|---|---|---|---|---|---|---|---|
| GOF | 44 | Marini et al. | p.Tyr411Cys | 14 yrs 8 m | M | paternally inherited | YES | Unclassified infantile epilepsy | moderate | 14 m | No | Learning disability, with some autistic traits | NR | normal |
| GOF /LOF | 45 | novel | p.Asp433Tyr | 3 yrs 8 m | M | de novo | YES | Infantile DEE | severe | 5.5 m | severe | No | NR | normal |
| wt-like (LOF in homo) | 46 | novel | p.Leu450Pro | 10 yrs 5 m | M | de novo | NO | | | | moderate | No | NR | normal |
| GOF | 47 | novel | p.Cys542Phe | died at 21 yrs | F | de novo | YES | Infantile DEE (West syndrome > LGS) | severe | 6m | severe | ASD, behavioural problems | Hypotonia, dysmorphisms wide based gait, sleep problems | dysgenesis of CC; abnormal sulcation of the frontal lobes |
| LOF | 48 | novel | p.Arg548His | 10 yrs | F | maternally inherited (with sz) | YES | Early onset absence epilepsy | moderate | 14 m | moderate | ADHD; severe learning disability LD; absent speech | large head, broad nasal base, poor motor coordination | normal |
| LOF | 49 | Marini et al. | p.Arg590Gln | 15 yrs | M | de novo | YES | CAE | mild | 72 m | No | NA | NR | normal |

LOF = loss-of-function, GOF = gain-of-function, PT = patient, AA = amino acid, DD = developmental delay, ASD = autism spectrum disorder, ADHD = attention deficit hyperactivity disorder, sz = seizures, yrs = years, m = months, F = female, M = male, NA = not available, NR = not reported, No = not present, GGE = generalized genetic epilepsy, DEE = developmental and epileptic encephalopathy, DS = Dravet syndrome, TCS = tonic-clonic seizure, FS+ = febrile seizures plus, DSA = specific learning disability, LD = learning disability, CAE = childhood absence epilepsy, LGS = Lennox-Gastaut syndrome, GEFS+ = genetic epilepsy with febrile seizures plus, [§]not included in clinical analysis
